## Supplementary Material for "The rise of grasslands is linked to atmospheric CO_2_ decline in the late Palaeogene"

#### Supporting Information

Luis Palazzesi[1,2,*], Oriane Hidalgo[2,3], Viviana D. Barreda[1], Félix Forest[2] and Sebastian Höhna[4,5]

[1]*Museo Argentino de Ciencias Naturales & Consejo Nacional de Investigaciones Científicas y Técnicas (CONICET), Buenos Aires C1405DJR, Argentina*
[2]*Jodrell Laboratory, Royal Botanic Gardens, Kew, Richmond, Surrey TW9 3DS, United Kingdom*
[3]*Institut Botànic de Barcelona (IBB, CSIC-Ajuntament de Barcelona), Catalonia, Spain*
[4]*GeoBio-Center, Ludwig-Maximilian-Universität München, 80333 Munich, Germany*
[5]*Department of Earth and Environmental Sciences, Paleontology & Geobiology, Ludwig-Maximilian-Universität München, 80333 Munich, Germany*

Correspondence should be sent to:
∗
∗∗

#### S1 Taxonomic representativeness of the most important vascular plant families in grasslands

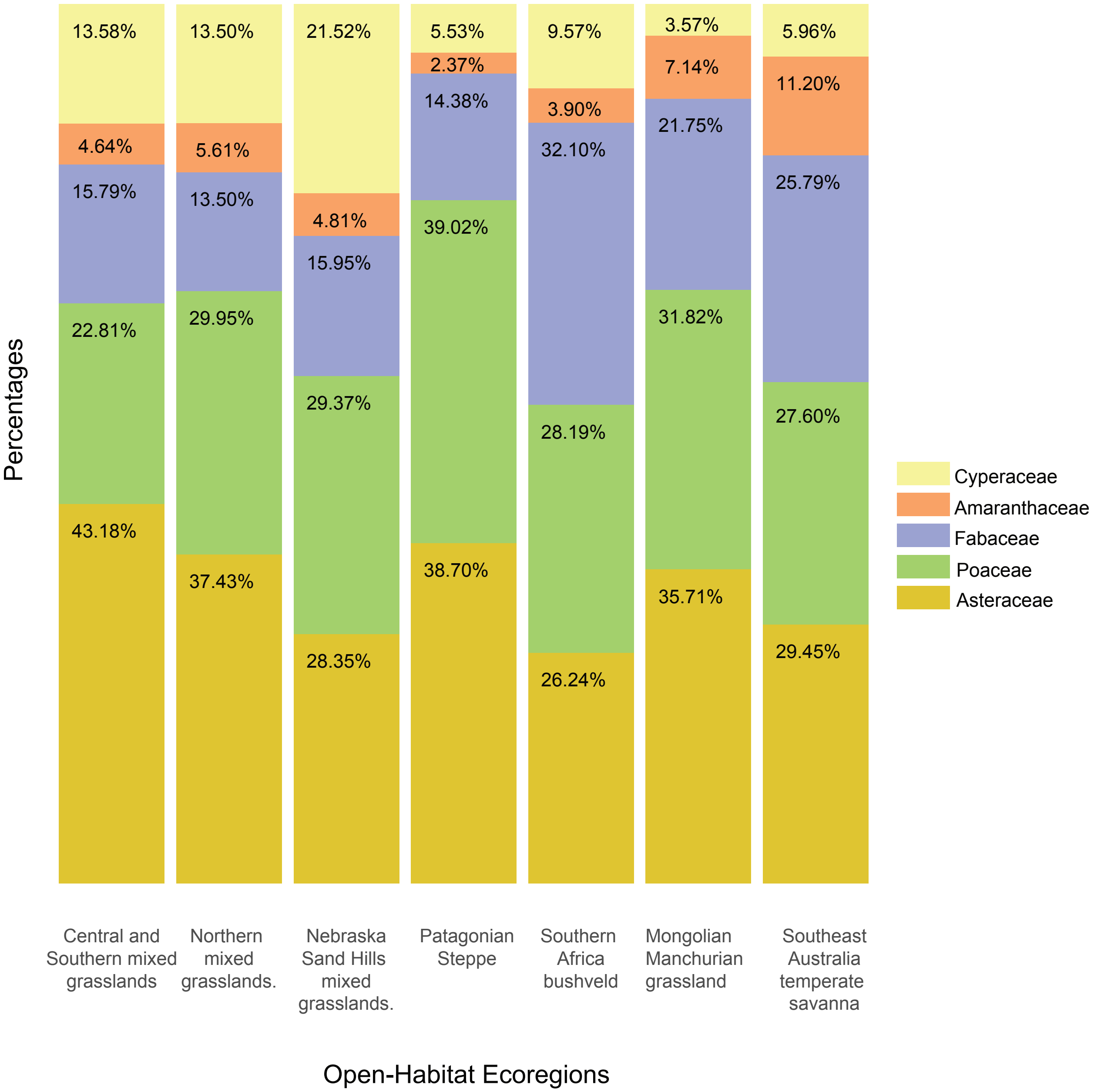

**Figure S1: Taxonomic representativeness of the most important vascular plant families in grasslands.** We selected seven WWF open-habitat eco-regions and quantified the number of species of the most abundant families using the GBIF portal through the R package 'rgbif' [5].

#### S2 Extent of major biomes (107 km$^2$) using an equilibrium vegetation model (BIOME4)

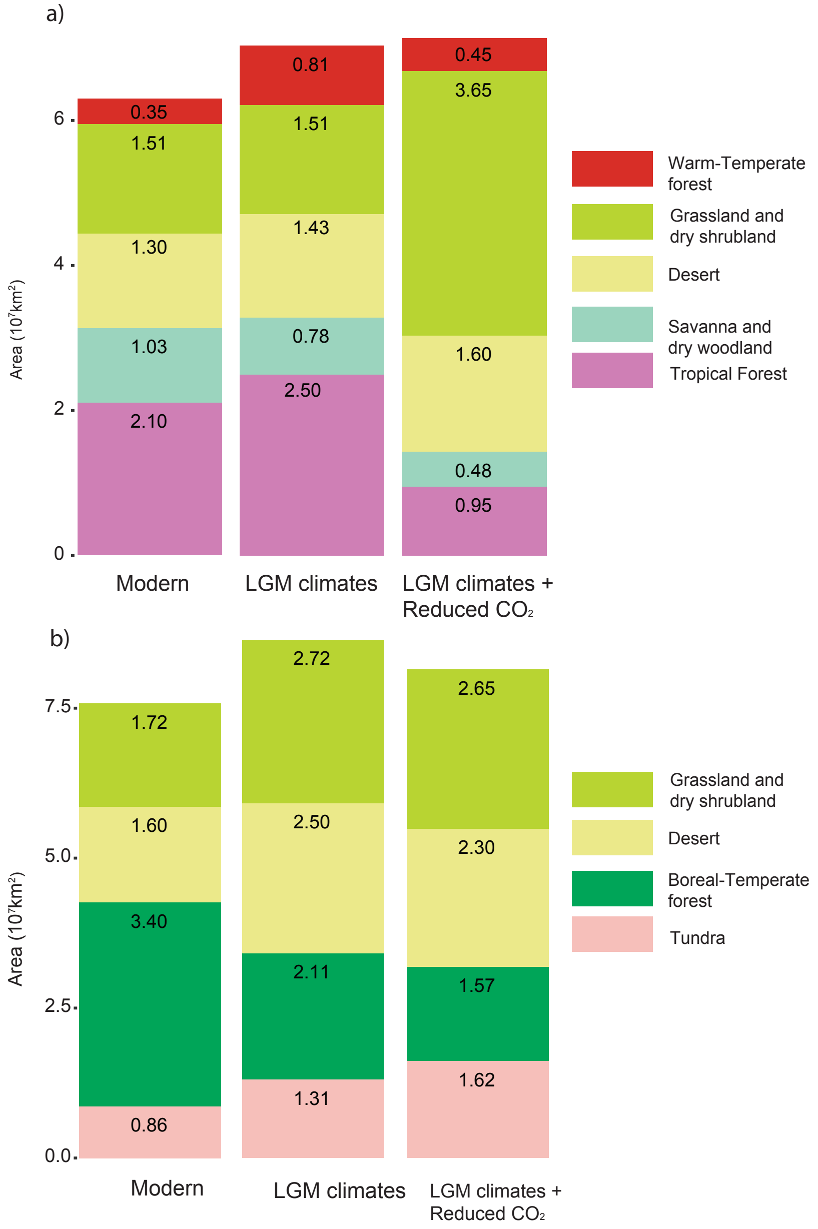

**Figure S2: Extent of major biomes (107 km$^2$) using an equilibrium vegetation model (BIOME4) [9].** a) Intertropical (30°N-30°S) extent of tropical forest, savanna and dry woodland, desert, grassland and dry shrubland and warm-temperate forest as a result of simulated LGM climate changes alone, LGM climate and reduced $CO_2$ concentrations compared to modern extent of the same biomes. b) Northern hemisphere extent of tundra, boreal-temperate forest, desert and grassland and dry shrubland as a result of simulated LGM climate changes alone, LGM climate plus reduced $CO_2$ conditions. The output of each of the seventeen climate simulations was averaged for each major biome in order to obtain a more compelling story about the effects of climate and $CO_2$ concentrations on the vegetation. Overall, the modeled physiological impact of low $CO_2$ produces i) a pronounced shift towards open-habitat vegetation, and ii) a reduction of ~50% of reduction in the area of tropical forests compared to the area simulated under modern climate and $CO_2$ conditions [9].

### S3 Asteraceae fossil species

**Table S1: Asteraceae fossil species.** Fossil data from Asteraceae come almost exclusively from pollen grains. The oldest Asteracean-like pollen comes from the latest Cretaceous of Antarctica and New Zealand [3], assigned to the Barnadesioideae, the sister subfamily to the core-Asteraceae. The unique macrofossil (inflorescence and associated pollen grains) confidently assigned to Asteraceae comes from the Middle Eocene of Patagonia showing similarities with the Mutisioideae and Carduoideae [2]. The earliest record of the Asteroideae (the clade that includes the most common open-habitat daisy tribes) occurs since the Late Oligocene of New Zealand but in very low frequencies. Fossils refer to this subfamily increased in abundance and diversity during the Miocene and Pliocene. Pollen referred to *Artemisia*, in particular, did not become abundant until the Middle-Late Miocene with several reports from central Europe, Asia and North America. Pre-Miocene findings need further verification. Overall, the Late Oligocene and in particular the Miocene witnessed the major step on the diversification of Asteraceae; ca. 80% of the fossil species recorded have been defined for this time interval.

| Species | Botanical Affinity | Epoch |
|---|---|---|
| *Tubulifloridites lilliei* type A *sensu* Barreda et al [3] | Barnadesioideae | Maastrichtian |
| *Tubulifloridites* sp. *sensu* Raine 2008 [15] | Mutisioideae/Carduoideae? | Early Eocene |
| *Mutisiapollis tellerieae* Barreda and Palazzesi | Mutisioideae/Carduoideae | Middle Eocene |
| *Raiguenrayun cura* Barreda et al | Mutisioideaea/Carduoideae | Middle Eocene |
| *Mutisiapollis patersonii* Macphail and Hill | Mutisioideae (Mutisieae, *Mutisia* type) | Early Oligocene |
| *Tubulifloridites antipodica* Cookson | Asteroideae | Oligocene/Miocene |
| *Tricolporopollenites microechinatus* Trevisan | Asteroideae | Oligocene/Miocene |
| *Mutisiapollis viteauensis* Barreda | Gochnatioideae (*Cnicothamnus* type) | Early Miocene |
| *Huanilipollis cabrerii* Barreda and Palazzesi | Mutisioideae (Nassauvieae) | Early Miocene |
| *Huanilipollis criscii* Barreda and Palazzesi | Mutisioideae (Nassauvieae) | Early Miocene |
| *Cichorieacidites ixeriformis* Zheng | Cichorioideae | Middle Miocene |
| *Lapsana* type Lancucka-Srodoniowa | Cichorioideae (Cichorieae) | Middle Miocene |
| *Tubulifloridites granulosus* Nagy | Asteroideae | Miocene |
| *Tubulifloridites simplis* Martin | Asteroideae | Miocene |
| *Echitricolporites spinosus* Germeraad et al | Asteroideae | Miocene |
| *Tubulifloridites anthemidearum* Nagy | Asteroideae (Anthemideae) | Miocene |
| *Artemisia* type *sensu* Leopold [13] | Asteroideae (Anthemideae) | Miocene |
| *Tubulifloridites ambrosinae* Nagy | Asteroideae (Ambrosiinae) | Miocene |
| *Tubulifloridites* sp. (*Ambrosia* type) Guler et al | Asteroideae (Heliantheae) | Miocene |
| *Xanthium* type Cavallo and Martinetto | Asteroideae (Heliantheae) | Miocene |
| *Tubulifloridites grandis* Nagy | Asteroideae? | Miocene |
| Compositae indet *sensu* Graham [8] | Asteroideae? | Miocene |
| *Quilembaypollis gamerroi* Palazzesi and Barreda | Barnadesioideae (*Chuquiraga type*) | Miocene |
| *Quilembaypollis tayuoides* Barreda and Palazzesi | Barnadesioideae (*Dasyphyllum* type) | Miocene |
| *Quilembaypollis stuessyi* Palazzesi and Barreda | Barnadesioideae (*Schlechtendalia* type) | Miocene |
| *Cirsium* type Lancucka-Srodoniowa | Carduoideae | Miocene |
| *Cichoriaearumpollenites gracilis* Nagy | Cichorioideae | Miocene |
| *Cichorium intybus* type Hochuli | Cichorioideae (Cichorieae) | Miocene |
| *Mutisiapollis* sp. *sensu* Barreda et al 2006 [1] | Mutisioideae (Mutisieae) | Miocene |
| *Tricolporopollenites microspinulitegillatus* Trevisan | Asteroideae (Anthemideae, *Artemisia* type) | Miocene/Pliocene |
| *Tricolporopollenites rarispinulitegillatus* Trevisan | Asteroideae (Anthemideae, *Artemisia* type) | Miocene/Pliocene |
| *Artemisiaepollenites minor* Zhu | Asteroideae (Anthemideae) | Miocene/Pliocene |
| *Tubulifloridites minutus* Regali | Asteroideae (Astereae, *Solidago* type) | Miocene/Pliocene |
| *Echitricolporites mcneillyi* Germeraad et al | Asteroideae (Heliantheae, Ambrosiinae) | Miocene/Pliocene |
| *Echitricolporites* sp. *sensu* Wang [18] | Asteroideae | Miocene/Pliocene |
| *Tubulifloridites* sp. *sensu* Wang [18] | Asteroideae? | Miocene/Pliocene |
| *Tricolporopollenites kozaniensis* Weiland | Carduoideae? | Miocene/Pliocene |
| *Tubulifloridites macroechinatus* Nagy | Carduoideae? | Miocene/Pliocene |
| *ricolporopollenites spinilophatus* Trevisan | Cichorioideae (Cichorieae, *Sonchus type*) | Miocene/Pliocene |
| *Cichorium* type *sensu* Blackmore [4] | Cichorioideae (Cichorieae) | Miocene/Pliocene |
| *Fenestrites longispinosus* Lorente | Cichorioideae (Cichorieae) | Miocene/Pliocene |
| *Liguliflorídites* sp. *sensu* Couper [7] | Cichorioideae (Cichorieae) | Miocene/Pliocene |
| *Sonchus* type *sensu* Blackmore [4] | Cichorioideae (Cichorieae) | Miocene/Pliocene |
| *Fenestrites spinosus* Hammen | Cichoroideae (Cichorieae/Vernoniae) | Miocene/Pliocene |
| *Artemisiaepollenites sellularis* Nagy | Asteroideae (Anthemideae) | Miocene/Pliocene |
| *Artemisiaepollenites leatus* Zheng | Asteroideae (Anthemideae) | Pliocene |
| *Scorzonera* type *sensu* Blackmore [4] | Cichorioideae (Cichorieae) | Pliocene |
| *Tubulifloridites pleistocenicus* Martin | Asteroideae | Pliocene/Pleistocene |

### S4 Asteraceae genes

**Table S2: Asteraceae genes** Number of taxa (total and sampled) and genes used in phylogenetic analyses for each clade of Asteraceae.

| Clades | Number of Species | Sampled Species | Age | Genes |
|---|---|---|---|---|
| Barnadesioideae | 92 | 70 | 64.41 | ITS, trnL, matK except 7 species of the backbone that include 12 markers (see Panero et al. 2015) |
| Famatinanthoideae | 1 | 1 | 60.82 | accD, atpB, matK, ndhD, ndhF, ndhI, ndhJKC, rbcL, rpoB, rpoC1exon2, trnL intron-trnL-F spacer |
| Mutisioideae | 637 | 210 | 40.55 | ITS, ndhF except 12 species of the backbone that include 12 markers |
| Stifftioideae | 35 | 4 | 34.84 | accD, atpB, matK, ndhD, ndhF, ndhI, ndhJKC, rbcL, rpoB, rpoC1exon1, rpoC1exon2, trnL intron-trnL-F spacer |
| Wunderlichioideae | 48 | 6 | 48.88 | accD, atpB, matK, ndhD, ndhF, ndhI, ndhJKC, rbcL, rpoB, rpoC1exon1, rpoC1exon2, trnL intron-trnL-F spacer |
| Gochnatioideae | 85 | 32 | 42.60 | ndhF, trnL except 3 species of the backbone that include 12 markers |
| Hecastocleidoideae | 1 | 1 | 46.86 | accD, atpB, matK, ndhD, ndhF, ndhI, ndhJKC, rbcL, rpoB, rpoC1exon1, rpoC1exon2, trnL intron-trnL-F spacer |
| Carduoideae | 2865 | 373 | 24.26 | psbA-trnH, matK except 5 species of the backbone that include 12 markers |
| Gymnarrhenoideae | 2 | 1 | 38.19 | accD, atpB, matK, ndhD, ndhF, ndhI, ndhJKC, rbcL, rpoB, rpoC1exon1, rpoC1exon2, trnL intron-trnL-F spacer |
| Corymbioideae | 7 | 1 | 32.13 | accD, atpB, matK, ndhD, ndhF, ndhI, ndhJKC, rbcL, rpoB, rpoC1exon1, rpoC1exon2, trnL intron-trnL-F spacer |
| Inuleae | 680 | 105 | 20.03 | psbA-trnH, trnL-F except 1 species of the backbone that include 12 markers |
| Helianthodae | 6611 | 510 | 21.41 | ITS, matK except 6 species of the backbone that include 12 markers |
| Senecionodae-Asterodae | 10589 | 648 | 26.60 | ITS, trnL-trnF except 3 species of the backbone that include 12 markers |
| Cichorioideae | 3994 | 759 | 28.51 | ITS, trnL-trnF except 4 species of the backbone that include 12 markers |
| Pertyoideae | 70 | 2 | 17.35 | accD, atpB, matK, ndhD, ndhF, ndhI, ndhJKC, rbcL, rpoB, rpoC1exon1, rpoC1exon2, trnL intron-trnL-F spacer |
| Total | 25717 | 2723 | | |

### S5 Poaceae genes

**Table S3: Poaceae genes** Number of taxa (total and sampled) and genes used in phylogenetic analyses for each clade of Poaceae according to Spriggs et al (2014). Age estimations correspond to the calibration scenario 1 of Spriggs et al (2014)[16]

| Clades | Number of Species | Sampled Species | Age | Genes |
|---|---|---|---|---|
| Pharoideae | 12 | 1 | 57.80 | matK, ndhF, rbcL |
| Puelioideae | 11 | 1 | 53.98 | matK, ndhF, rbcL |
| Ehrhartoideae | 112 | 66 | 33.77 | GPA1, ITS, matK, ndhC, ndhF, psbHpetB, psbZ, rbcL rps19, trnHpsbA, ycf3 |
| Bambusoideae | 1441 | 418 | 23.27 | matK, ndhF, rbcL, ITS, psbA-trnH, rpL16, rpL32, rps16-trnQ, trnC-rpoB, trnL-trnF, trnTtrnD |
| Pooideae | 3850 | 1335 | 38.25 | DMC1, ITS, matK, ndhF, pgk1, psbAtrnH, rbcL, rpb2, rpoA, rps19, trnKrps16, trnLtrnF |
| Aristidoideae | 365 | 125 | 27.95 | ITS, matK, ndhF, rbcL, rpL16, trnL-trnF |
| Centotheceae | 33 | 33 | 29.66 | ndhF, rbcL, matk, trnLtrnF, phyB, ITS |
| Andropogoneae | 1274 | 250 | 16.97 | ITS, phyB, trnLtrnF, matK, ndhF, rbcL |
| Paspaleae | 664 | 168 | 21.01 | trnL-trnF, rbcL, psbA-trnH, atpB-rbcL, trnG, rpL16, ndhF, matK, ITS |
| Paniceae-Gynerieae | 1254 | 387 | 21.29 | ITS, kn1, matK, ndhF, rbcL, rpL16, trnLtrnF |
| Danthonioideae | 281 | 234 | 28.71 | ndhF, rbcL, atpB-rbcL, ITS, psbM, rpl16, rpoC2, trnC-trnD, trnL, ycf6-trnC, trnT-trnL, matK |
| Chloridoideae | 1721 | 534 | 32.91 | ITS, matK, ndhF, rbcL rl16 rps16, rps3, trnL-trnF |
| Arundinoideae-Micrairoideae | 188+46 | 43 | 30.38 | ITS, rpoC2, matK, ndhF, rbcL |
| Total | 11256 | 3595 | | |

#### S6 Environmental $CO_2$ curve

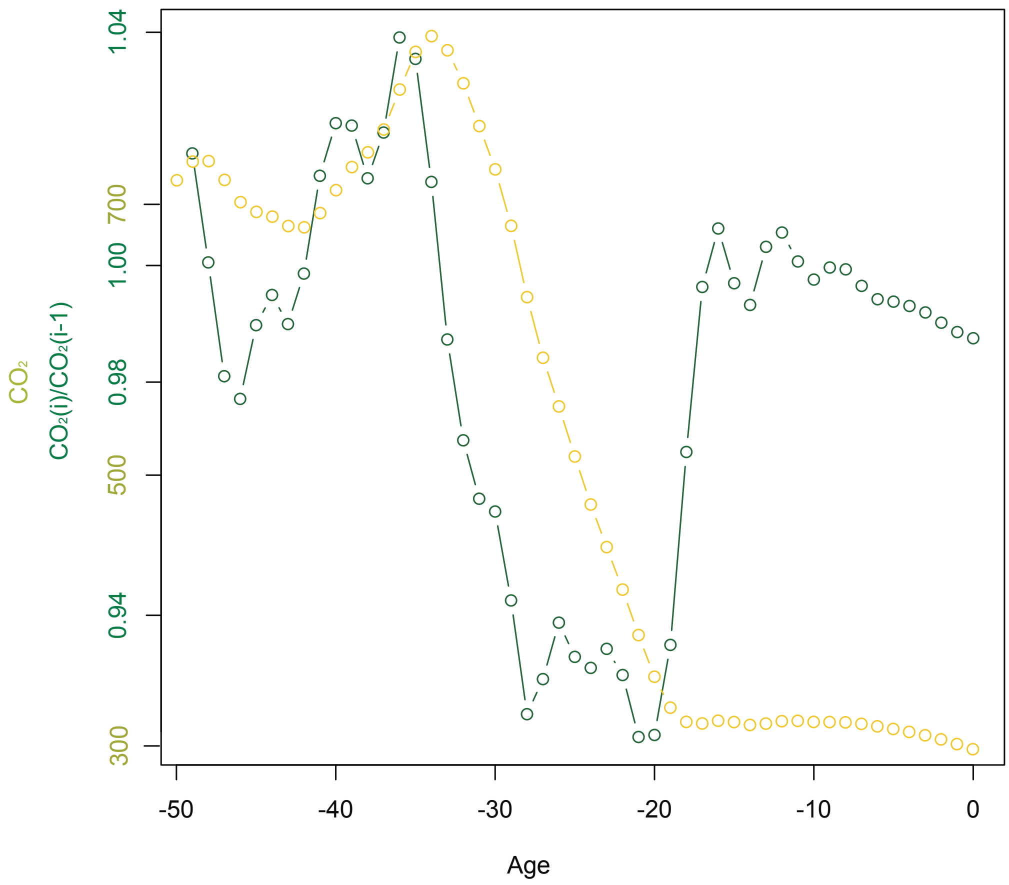

**Figure S3: Environmental $CO_2$ and changes in $CO_2$ between time intervals**. The lighter green line shows the environmental $CO_2$ as was used in the environmentally-dependent diversification rates models. The darker green line shows the changes in $CO_2$ over time, which might translate into changes in diversification rates if diversification rates are correlated with environmental $CO_2$.

### S7 Models and Priors for the environmental correlation analyses

#### S7.1 Fixed

Our first model, which is equivalent to the exponential environmentally-dependent diversification model of Condamine et al. [6], has two parameters per speciation and extinction rate; the initial rate $\lambda_0$ and $\mu_0$ and the correlation coefficient $\beta$, with

$$\lambda_0 \sim \text{Uniform}(0, 100) \tag{S1}$$
$$\ln(\lambda_i) = \ln(\lambda_{i-1}) + \beta_\lambda \times \Delta CO_2 \quad . \tag{S2}$$

**Table S4: Model parameter names and prior distributions for the fixed model.**

| Parameter | $X$ | $f(X)$ |
|---|---|---|
| Speciation at present | $\lambda_0$ | Uniform(0,100) |
| Extinction at present | $\mu_0$ | Uniform(0,100) |
| Speciation Correlation | $\beta_\lambda$ | Normal(0,1) |
| Extinction Correlation | $\beta_\mu$ | Normal(0,1) |

## S7.2 UC

The second models extends our first model by addition additional "errors", or variation, on top of the diversification rate variation due to the environmental variable. Thus, we additionally have a variance parameter $\sigma^2$ as well as $\epsilon_\lambda$ and $\epsilon_\mu$ per interval. The resulting speciation and extinction rates are computed by

$$\lambda_0 \sim \text{Uniform}(0, 100) \tag{S3}$$
$$\ln(\hat{\lambda}_i) = \ln(\hat{\lambda}_{i-1}) + \beta_\lambda \times \Delta CO_2 \tag{S4}$$
$$\epsilon_i \sim \text{Normal}(0, \sigma) \tag{S5}$$
$$\ln(\lambda_i) = \ln(\hat{\lambda}_i) + \epsilon_i \quad . \tag{S6}$$

**Table S5: Model parameter names and prior distributions for the UC model.**

| Parameter | $X$ | $f(X)$ |
|---|---|---|
| Speciation at present | $\lambda_0$ | Uniform(0,100) |
| Extinction at present | $\mu_0$ | Uniform(0,100) |
| Speciation Variation | $\sigma_\lambda$ | HalfCauchy(0,1) |
| Extinction Variation | $\sigma_\mu$ | HalfCauchy(0,1) |
| Speciation Correlation | $\beta_\lambda$ | Normal(0,1) |
| Extinction Correlation | $\beta_\mu$ | Normal(0,1) |
| Per interval log-speciation rate variation | $\epsilon_\lambda$ | Normal(0,$\sigma_\lambda$) |
| Per interval log-extinction rate variation | $\epsilon_\mu$ | Normal(0,$\sigma_\mu$) |

#### S7.3 Gaussian Markov Random Field (GMRF)

The GMRF model has autocorrelated rate variation compared to the uncorrelated variation from the UC model. The speciation and extinction rates are computed by

$$\lambda_0 \sim \text{Uniform}(0, 100) \quad \text{(S7)}$$
$$\sigma \sim \text{halfCauchy}(0, 1) \quad \text{(S8)}$$
$$\ln(\lambda_i) \sim \text{Normal}(\ln(\lambda_{i-1}) + \beta_\lambda \times \Delta CO_2), \sigma\zeta) \quad . \quad \text{(S9)}$$

We additionally used a global scaling parameter for the variation of diversification of $\zeta = 0.587405 \times N$, where $N$ is the number of epochs, so that we expect one order of magnitude variation over the course of the age of the phylogeny [14]. Alternatively, the parameter $\zeta$ could be place into the prior distribution on $\sigma$ by using a halfCauchy$(0, \zeta)$ prior distribution instead. Both approaches are equivalent.

**Table S6: Model parameter names and prior distributions for the GMRF model.**

| Parameter | $X$ | $f(X)$ |
|---|---|---|
| Speciation at present | $\lambda_0$ | Uniform(0,100) |
| Extinction at present | $\mu_0$ | Uniform(0,100) |
| Speciation Variation | $\gamma_\lambda$ | HalfCauchy(0,1) |
| Extinction Variation | $\gamma_\mu$ | HalfCauchy(0,1) |
| Speciation Correlation | $\beta_\lambda$ | Normal(0,1) |
| Extinction Correlation | $\beta_\mu$ | Normal(0,1) |
| Per interval log-speciation rate difference | $\ln(\Delta_\lambda)$ | Normal(0,$\gamma_\lambda\zeta$) |
| Per interval log-extinction rate difference | $\ln(\Delta_\mu)$ | Normal(0,$\gamma_\mu\zeta$) |

#### S7.4 HSMRF

The HSMRF model allows for interval specific variance parameters $\gamma_i^2$. This allows for some epochs to be more variable, while most epochs will actually be less variable [14]. The resulting speciation and extinction rates are thus computed by

$$\lambda_0 \sim \text{Uniform}(0, 100) \quad \text{(S10)}$$
$$\sigma \sim \text{halfCauchy}(0, 1) \quad \text{(S11)}$$
$$\gamma_i \sim \text{halfCauchy}(0, 1) \quad \text{(S12)}$$
$$\ln(\lambda_i) \sim \text{Normal}(\ln(\lambda_{i-1}) + \beta_\lambda \times \Delta CO_2), \sigma\gamma_i\zeta) \quad . \quad \text{(S13)}$$

As before, we additionally used a global scaling parameter for the variation of diversification of $\zeta = 0.587405 \times N$ so that we expect one order of magnitude variation over the course of the age of the phylogeny [14].

**Table S7: Model parameter names and prior distributions for the HSMRF model.**

| Parameter | $X$ | $f(X)$ |
|---|---|---|
| Speciation at present | $\lambda_0$ | Uniform(0,100) |
| Extinction at present | $\mu_0$ | Uniform(0,100) |
| Speciation Variation | $\sigma_\lambda$ | HalfCauchy(0,1) |
| Extinction Variation | $\sigma_\mu$ | HalfCauchy(0,1) |
| Speciation Correlation | $\beta_\lambda$ | Normal(0,1) |
| Extinction Correlation | $\beta_\mu$ | Normal(0,1) |
| Per interval speciation rate variation factor | $\gamma_{\lambda,i}$ | HalfCauchy(0,1) |
| Per interval extinction rate variation factor | $\gamma_{\mu,i}$ | HalfCauchy(0,1) |
| Per interval log-speciation rate difference | $\ln(\Delta_{\lambda,i})$ | Normal(0,$\gamma_{\lambda,i}\sigma_\lambda\zeta$) |
| Per interval log-extinction rate difference | $\ln(\Delta_{\mu,i})$ | Normal(0,$\gamma_{\mu,i}\sigma_\mu\zeta$) |

#### S8 Diversification rates with phylogenies

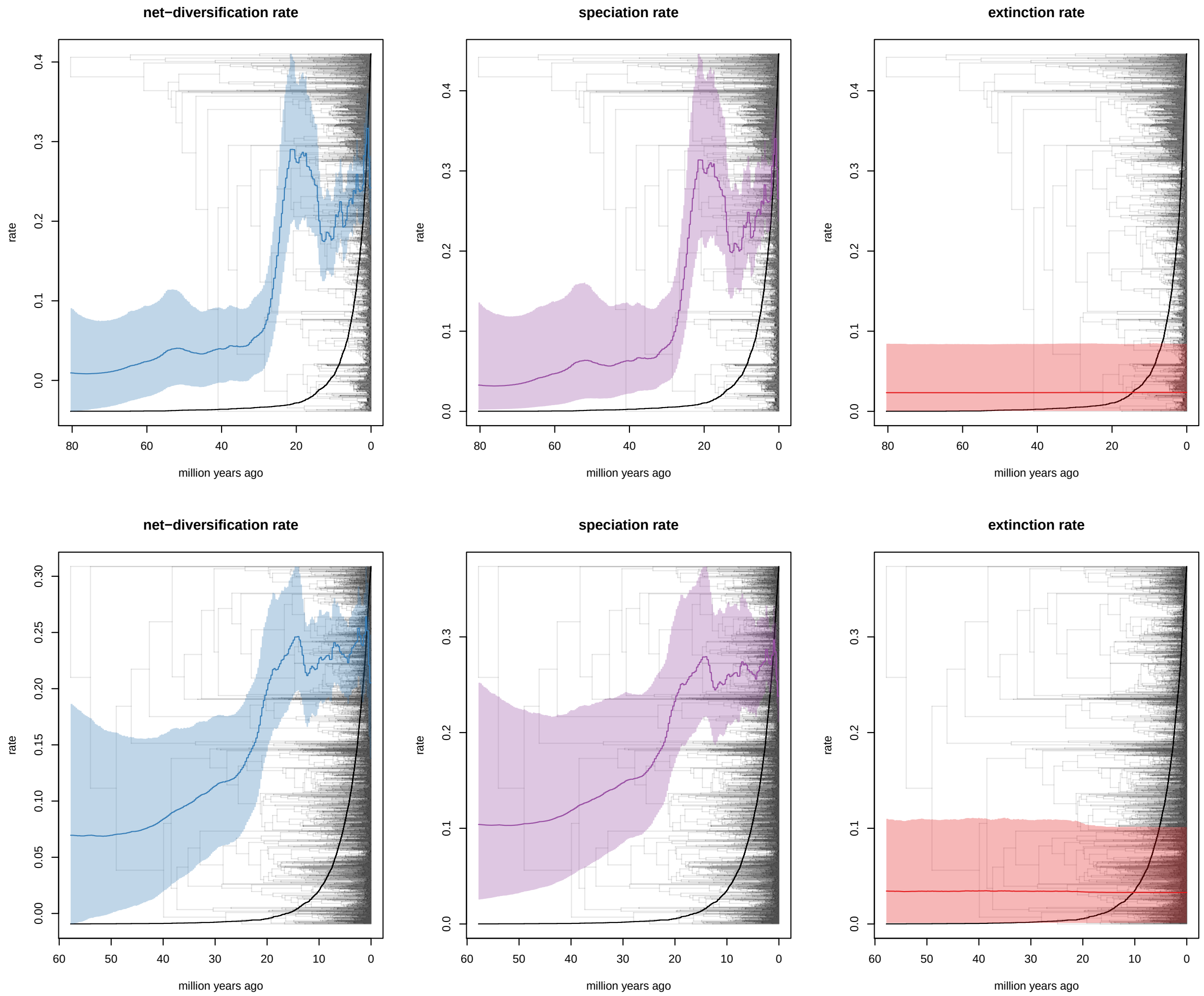

**Figure S4: Net-diversification, speciation and extinction rates with daisy (top) and grass (bottom) phylogenies.** Note that the most pronounced increase in net-diversification rates in both families occurred well before the divergence of most of the living species. This is because our *empirical* taxon sampling integrated missing (non-sampled) species within the most specious clades.

#### S9 The effect of different prior models on diversification rates

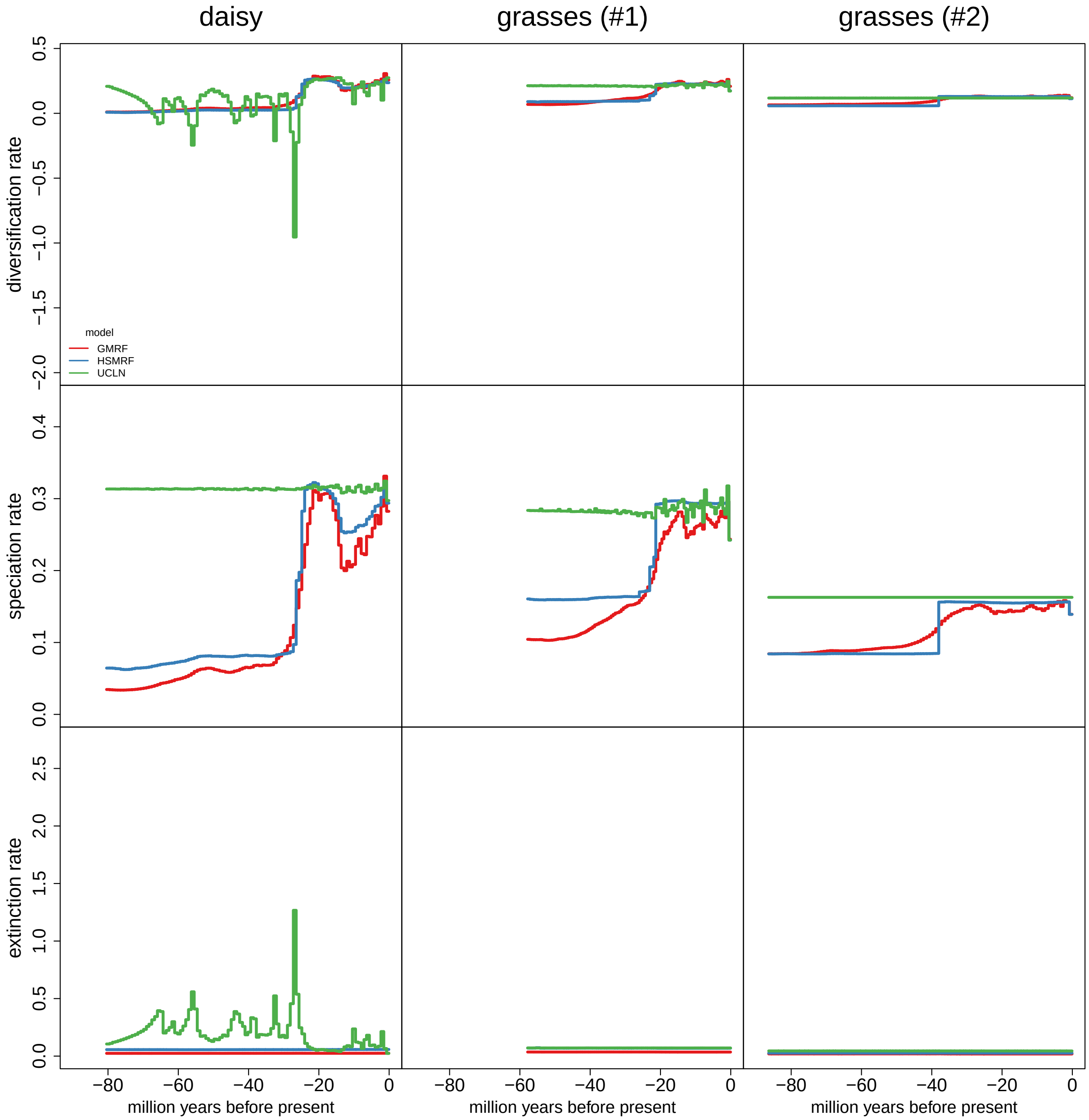

**Figure S5: Diversification rate estimates using different prior models.** The speciation, extinction and net-diversification rate estimates using the GMRF (red), HSRMF (blue) and UCLN (green) prior models. The rates were estimated using 100 epochs (for other numbers of epochs see Figure S6). The two autocorrelated models qualitatively agree on the estimated rates and infer diversification rate shifts driven by changes in the speciation rate. As expected, the GMRF model produces smoother rates while the HSMRF model produces more discrete jumps. The UCLN model produces strong variation which is a clear result of over-fitting (see Figure S7).

### S10 The effect of number of epochs on diversification rates

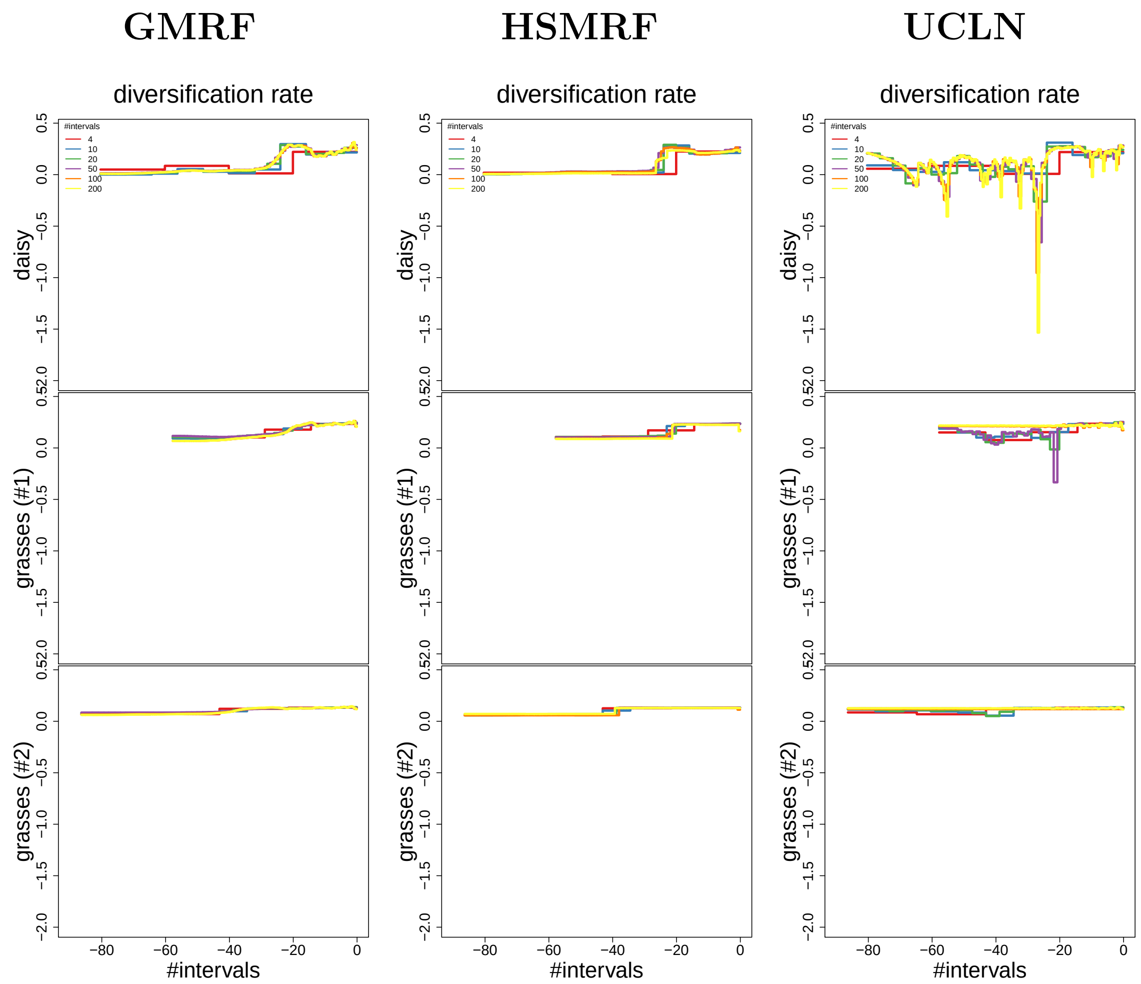

**Figure S6: Diversification rate estimates for different number of epochs.** We estimated diversification rates using $N = \{4, 10, 20, 50, 100, 200\}$ epochs for all three diversification prior models (GMRF, HSMRF and UCLN). The two autocorrelated models estimate smoother rate functions with higher number of intervals, as previously observed by Magee et al[14]. The uncorrelated model over-fitted when many intervals where used (see Figure S7).

#### S11 Model selection of diversification models

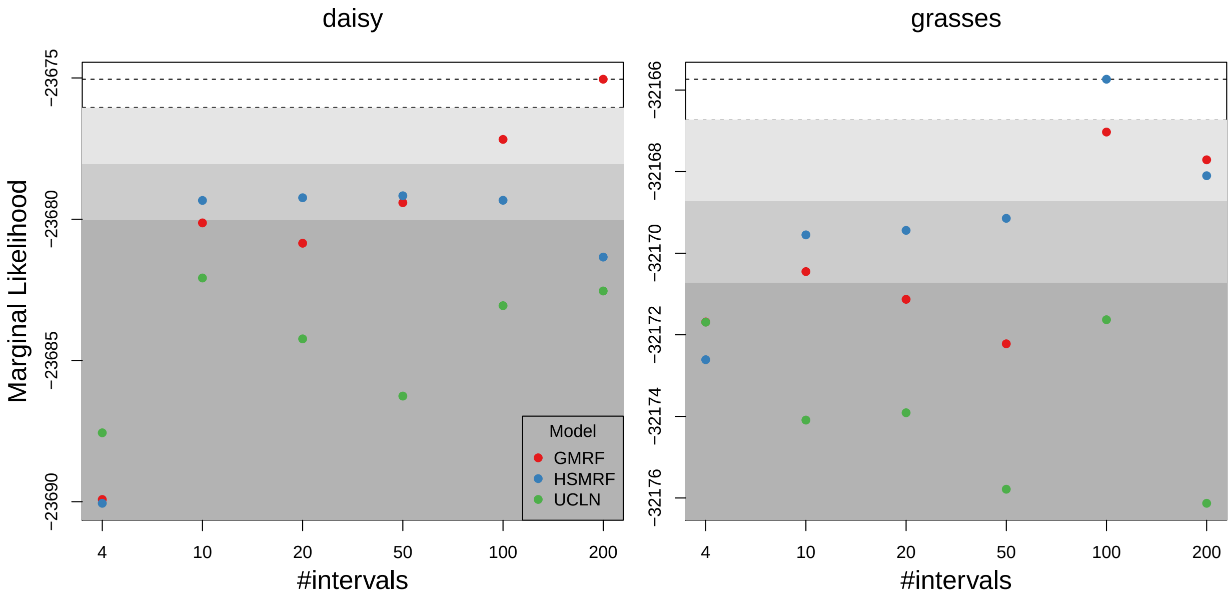

**Figure S7: Model selection between the three diversification rate prior models and for different number of epochs.** We estimated marginal likelihoods for all three diversification prior models (GMRF, HSMRF and UCLN) and for varying number of epochs, $N = \{4, 10, 20, 50, 100, 200\}$. The colored areas show (i) no significant support in white, (ii) support in light gray, (iii) strong support in gray, and (iv) decisive support in dark gray according to the common Bayes factor thresholds [12]. The UCLN model was always decisively rejected. The best fitting model for the daisy phylogeny was the GMRF with 200 epochs and the HSRMF with 100 epochs for the grasses phylogeny. Interestingly, only for $N = 4$ epochs was the UCLN model better (daisy dataset) or equially good (grasses dataset) but more epochs resulted in worse model fit (best fit for daisies was 10 epochs and 4 epochs for grasses).

#### S12 The effect of different taxon sampling methods

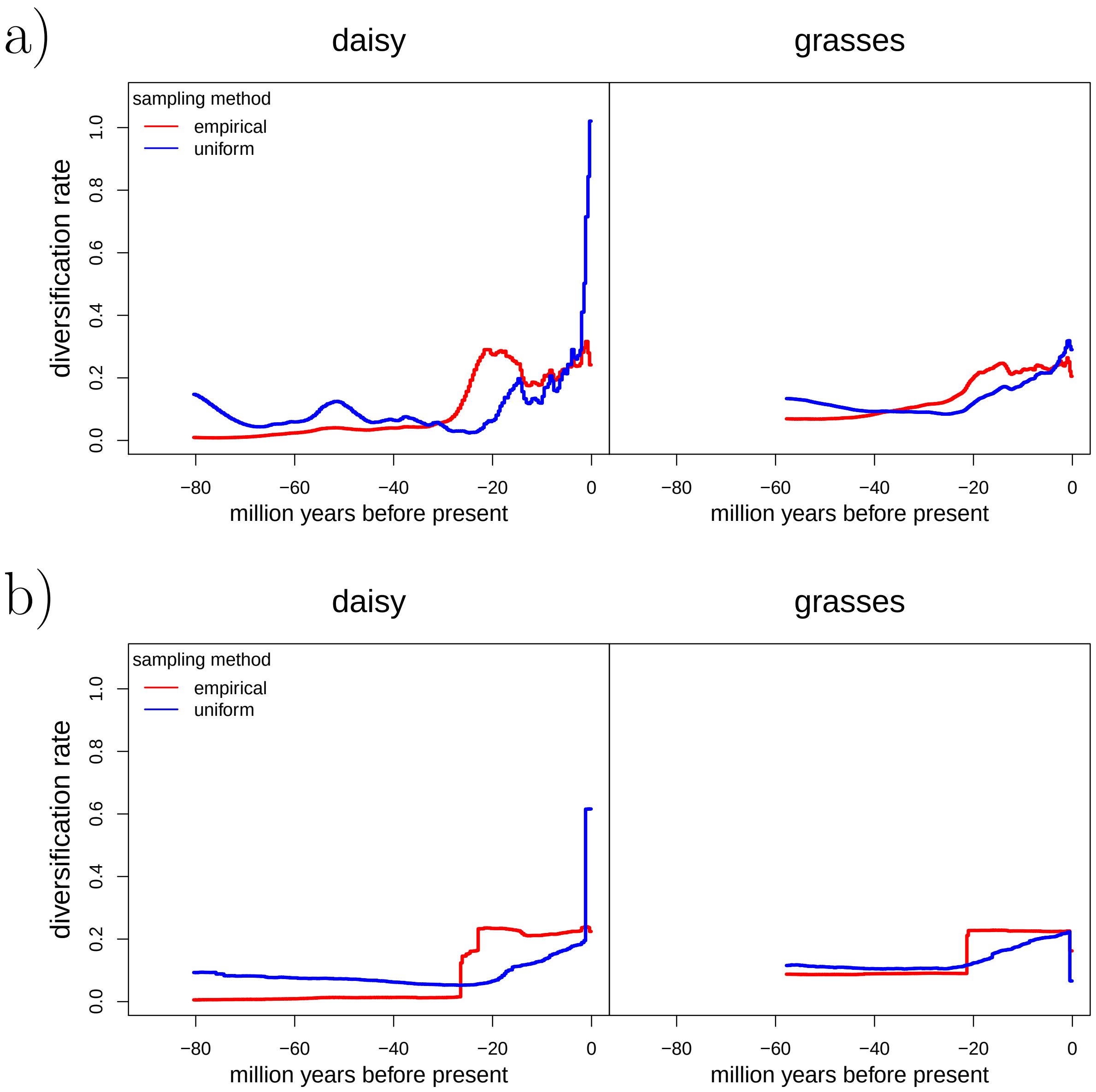

**Figure S8: Diversification rate estimates using different incomplete taxon sampling methods.** We estimated the diversification rates using our newly developed *empirical* taxon sampling and the previously developed *uniform* taxon sampling. The diversification rates were estimates using a) the GMRF diversification rate prior model and b) the HSRMF diversification rate prior model. For each model with assumed $N = 200$ epochs. The estimated diversification rate differ depending on the incomplete taxon sampling method, but are similar between diversification rate prior models (see Figure S5). Thus, using the correct incomplete taxon sampling approach is crucial for unbiased estimation of diversification rates.

### S13 The effect of different models on correlations to the environmental variable

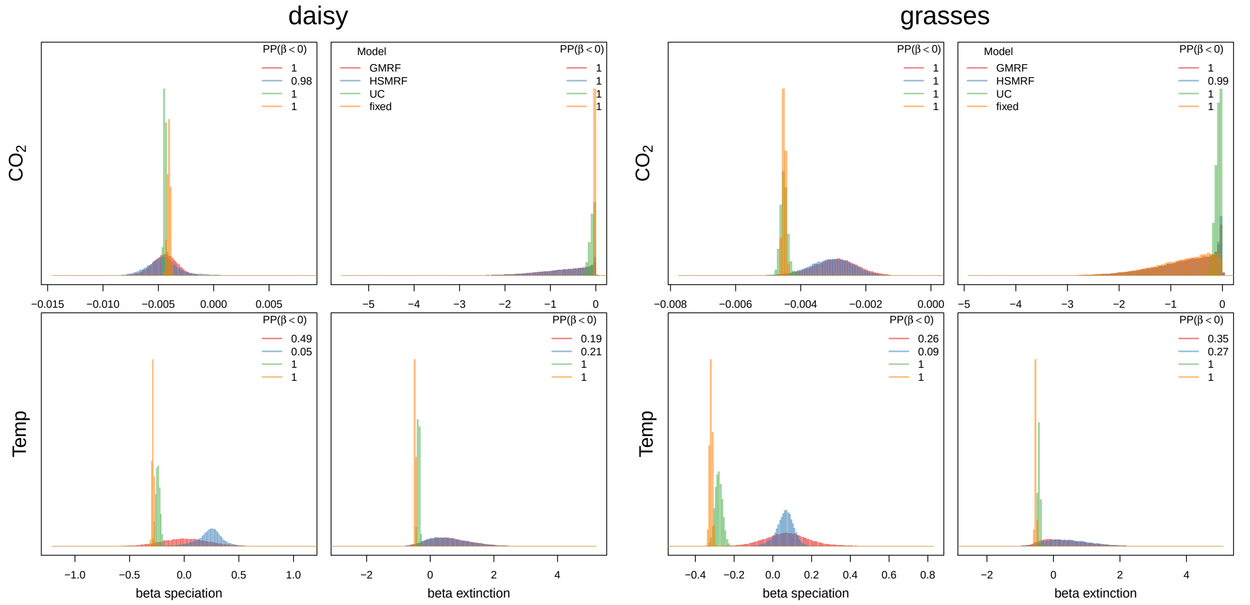

**Figure S9: Estimating correlation between diversification rates and two environmental variables.** We estimated the correlation between atmospheric $CO_2$ and paleo-temperature (with environmental variables averaged in 1MY bins) to speciation and extinction rates using our four environmentally-dependent diversification models: *fixed*, UC, GMRF and HSMRF. The inset posterior probabilities show the posterior probability that the correlation coefficient $\beta$ is smaller than 0 (i.e., a negative correlation). If the posterior probability is close to one, then there is a significant negative correlation and if the posterior probability is close to zero, then there is a significant positive correlation. The speciation rate is negatively correlated with atmospheric $CO_2$ for the both the daisy and grasses datasets regardless of the chosen model. The extinction rate is also negatively correlated and shows the same trend as the speciation rate. The correlation between paleo-temperature and speciation rate is less certain: the *fixed* and UC model infer a negative correlation whereas the GMRF and HSMRF infer no correlation or a positive correlation. The HSMRF model was the best fit model for the paleo-temperature analyses (see Figure S10). The same trend is seen in both datasets for the daisies and grasses.

### S14 Model selection of environmental variable correlated models

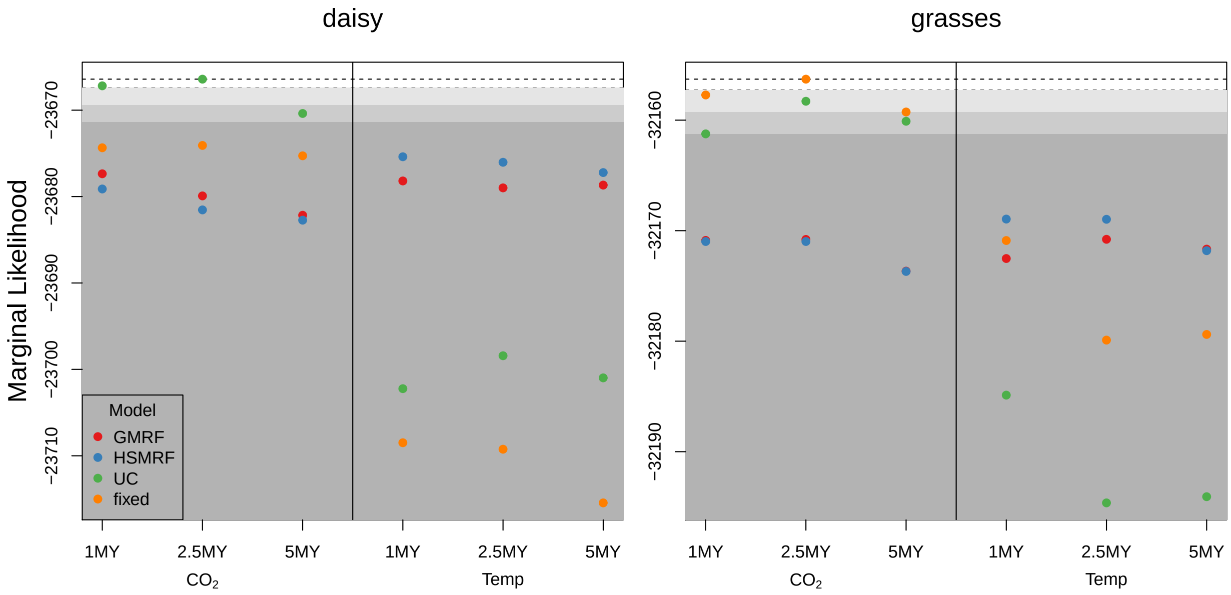

**Figure S10: Marginal likelihood estimates for the four environmentally-dependent diversification models.** We estimated marginal likelihoods using stepping stone sampling for the four environmentally-dependent diversification models (*fixed*, UC, GRMF and HSMRF) and two environmental variables (atmospheric $CO_2$ and paleo-temperature). Additionally, we binned the environmental variable in 1, 2.5 and 5 million year bins. We computed the marginal likelihoods for both the daisy and grasses dataset (calibration method #1). The shaded boxes represent significance levels according to standard Bayes factors[12]: slightly support (white), supported (light gray), strongly supported (dark gray) and decisively supported (dark gray). We found that for the daisy dataset the best predictor is atmospheric $CO_2$ with 2.5 MY bins using the UC model. The binning size did not impact the results and the order of preferred models remained unchanged. For the grasses dataset, the *fixed* model with atmospheric $CO_2$ as predictor variable in 2.5 MY bins received highest support. Interestingly, for the paleo-temperature as predictor variable the results flip with the two autocorrelated models (GMRF and HSMRF) receiving higher marginal likelihood than the *fixed* and UC model. This is particularly important as for the atmospheric $CO_2$ all models agreed on the negative correlation between $CO_2$ and speciation rates, whereas for the paleo-temperature the results qualitatively differed from negative correlation (*fixed* and UC) to no correlation or slighly positive correlation (GMRF and HSMRF); see Figure S9.

#### S15 The effect of binning on environmental variable correlated models

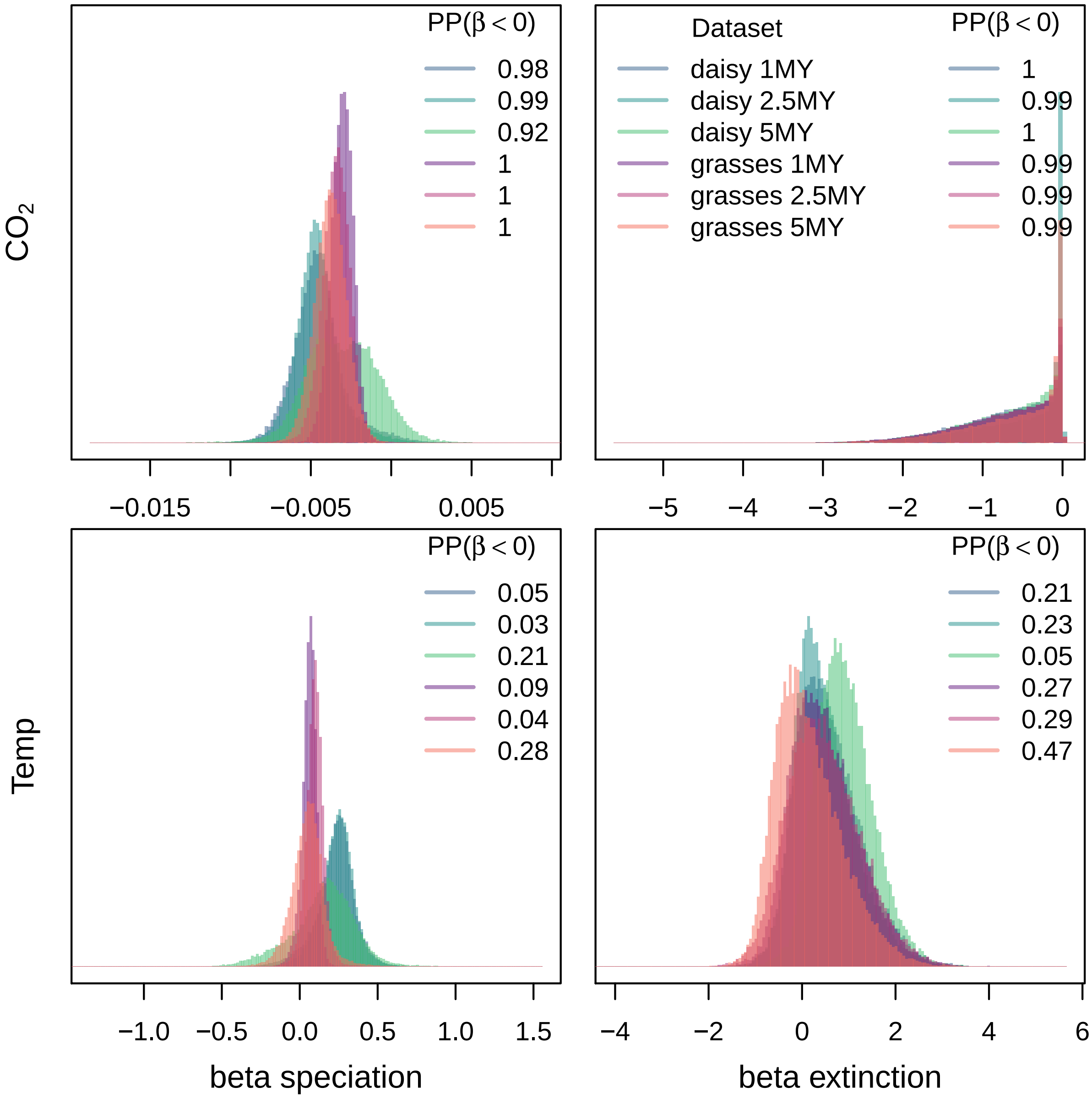

**Figure S11: Estimated correlation coefficient for different bin size of the environmental variable.** We estimated the correlation coefficient $\beta$ between the speciation and extinction rates and the environmental variable (atmospheric $CO_2$ and paloe-temperature) using the HSRMF model. The environmental variables were computed as averages for bins of size 1, 2.5 and 5 MY. The inset posterior probabilities show the posterior probability that the correlation coefficient $\beta$ is smaller than 0 (i.e., a negative correlation). If the posterior probability is close to one, then there is a significant negative correlation and if the posterior probability is close to zero, then there is a significant positive correlation. There is little effect on the correlation coefficient depending on the bin size, although a bin size of 5MY has the widest posterior distribution and is slightly shifted closer towards zero values and the posterior probabilities was less significant.

### S16 The effect of node calibration on environmental variable correlated models

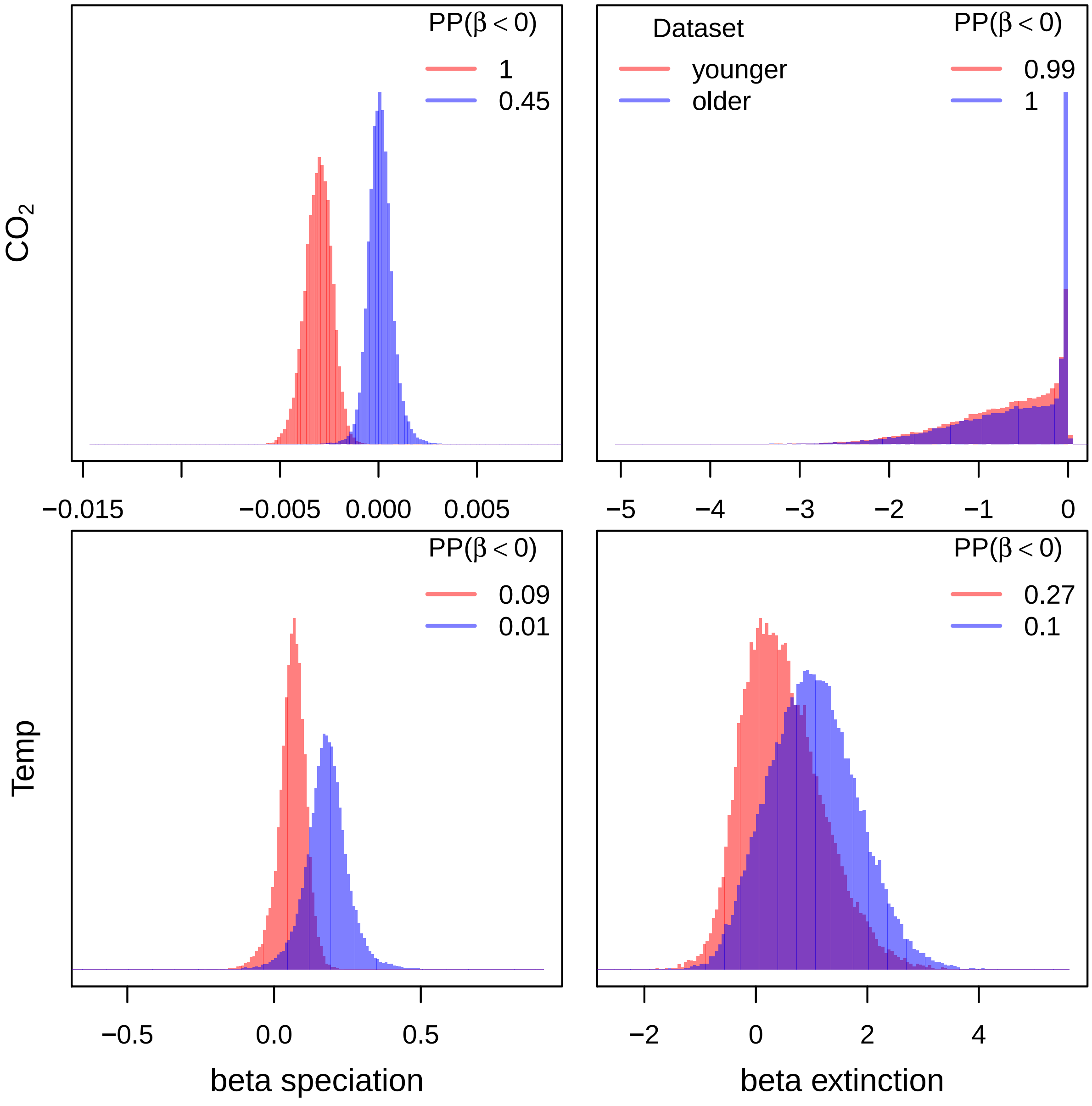

**Figure S12: Estimated correlation coefficient $\beta$ for the grasses dataset depending on node calibration scenario.** We compared the correlation coefficient between speciation and extinction rates and environmental variables (atmospheric $CO_2$ and paleo-temperature with averages over 1MY bins). The inset posterior probabilities show the posterior probability that the correlation coefficient $\beta$ is smaller than 0 (i.e., a negative correlation). If the posterior probability is close to one, then there is a significant negative correlation and if the posterior probability is close to zero, then there is a significant positive correlation. We applied the HSRMF model to the two grasses phylogenies obtained under different calibration scenarios (scenario #1 younger and scenario #2 older). For scenario #1 we estimate a negative correlation between atmospheric $CO_2$ and speciation rates but not correlation to paleo-temperature. For scenario #2 we estimate a positive correlation between speciation rates and paleo-temperature but no correlation to atmospheric $CO_2$.

### S17 The effect of environmentally-dependent diversification models on diversification rates

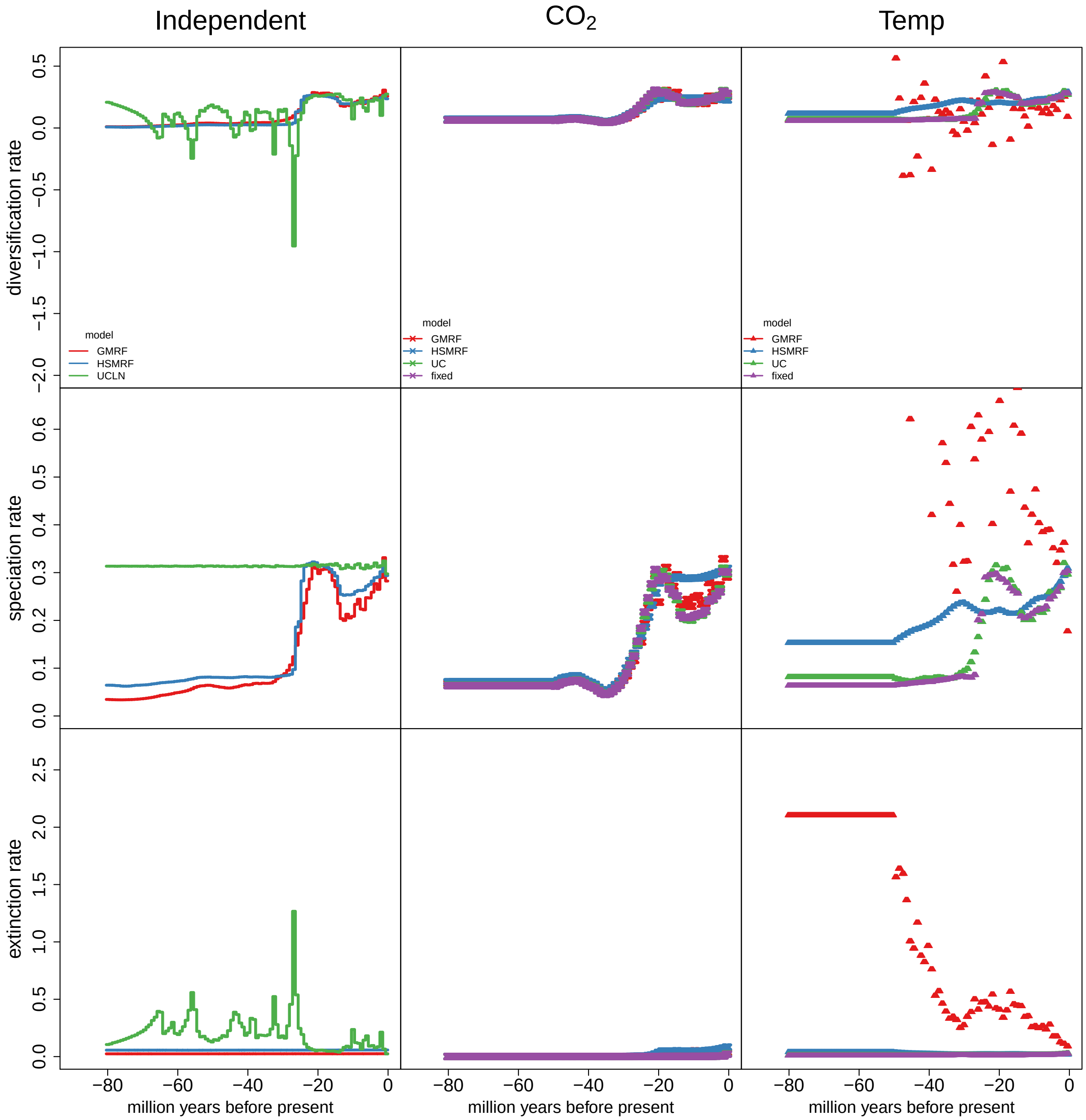

**Figure S13: Estimated diversification rates for the daisy phylogeny.** Here we compare the estimated diversification rates of our environmentally-dependent models to the null-hypothesis without environmental dependence. The environmentally dependent HSMRF and GMRF models yield extremely similar diversification rates compared with the environmentally independent diversification rate estimates. This shows that the diversification rates are not forced due the environmental correlation, but instead are driven by the information in the data (the phylogeny with divergence times). The uncorrelated (UC) rate models shows very different diversification rates for the correlation with paleo-temperature which confirms the disagreement in correlation coefficients (Figure S9). Unsurprisingly, the *fixed* environmentally-dependent diversification rate model show diversification rate closely following the environmental variable because no other source of rate-variation is allowed. Thus, all diversification rate variation in the *fixed* model, e.g., when using paleo-temperature as the environmental variable, is misleading because it is wrongfully enforced by the environmental variable.

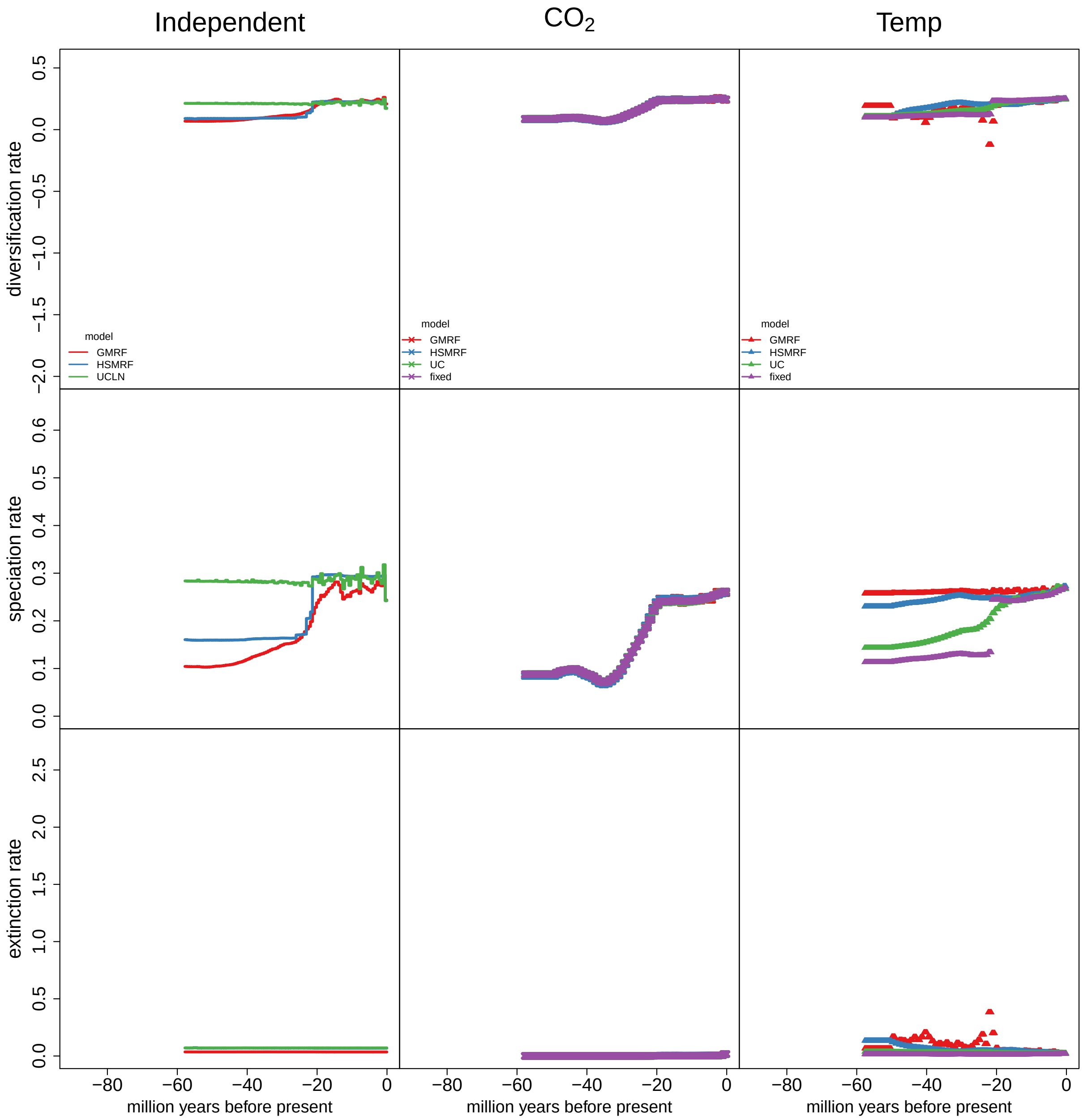

**Figure S14: Estimated diversification rates for the grasses phylogeny (calibration scenario 1).** Here we compare the estimated diversification rates of our environmentally-dependent models to the null-hypothesis without environmental dependence. For the $CO_2$ as the environmental driver, we see that all four environmentally-dependent diversification models provide very similar diversification rates. This confirms the agree that all models infer a correlation between $CO_2$ and diversification rates in grasses (Figure S9). Conversely, the diversification rates of all four models disagree if paleo-temperature is used. Thus, the correlation between paleo-temperature and diversification rates is unreliable and dependent on the specific model.

### S18 Simulation study of environmentally-dependent diversification rates

#### S18.1 Simulated diversification rates under environmentally-dependent diversification rates models

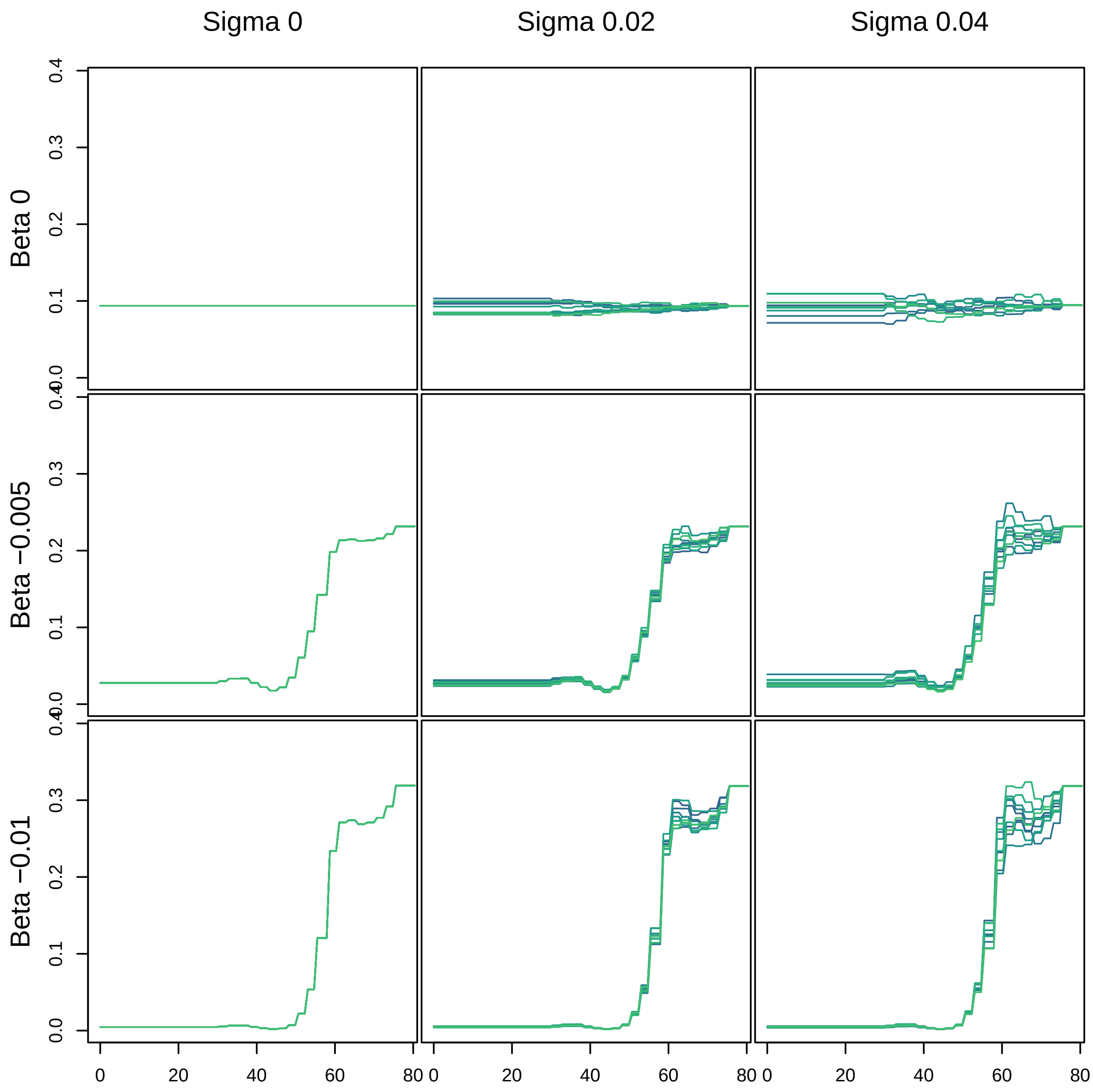

**Figure S15: Speciation rated simulated under an environmentally-dependent GRMF diversification model.** We simulated speciation rates with different correlation coefficients $\beta = \{0, -0.005, -.01\}$ and standard deviations $\sigma = \{0, 0.02, 0.04\}$. We used the atmospheric $CO_2$ as the environmental variable. When $\beta = 0$ in the simulations, then this model was equivalent to an environmentally-independent diversification model. When $\sigma = 0$ in the simulations, then this model was equal to the *fixed* environmentally-diversification model. When both $\beta = 0$ and $\sigma = 0$, then we simulated under a constant rate process (top left). This establishes the false positive rate in our simulations. Note that the autocorrelated model (GMRF) has overall more variation in speciation rates (e.g., top right) compared to the uncorrelated model (Figure S16) because the variation is additive over epochs in the autocorrelated model.

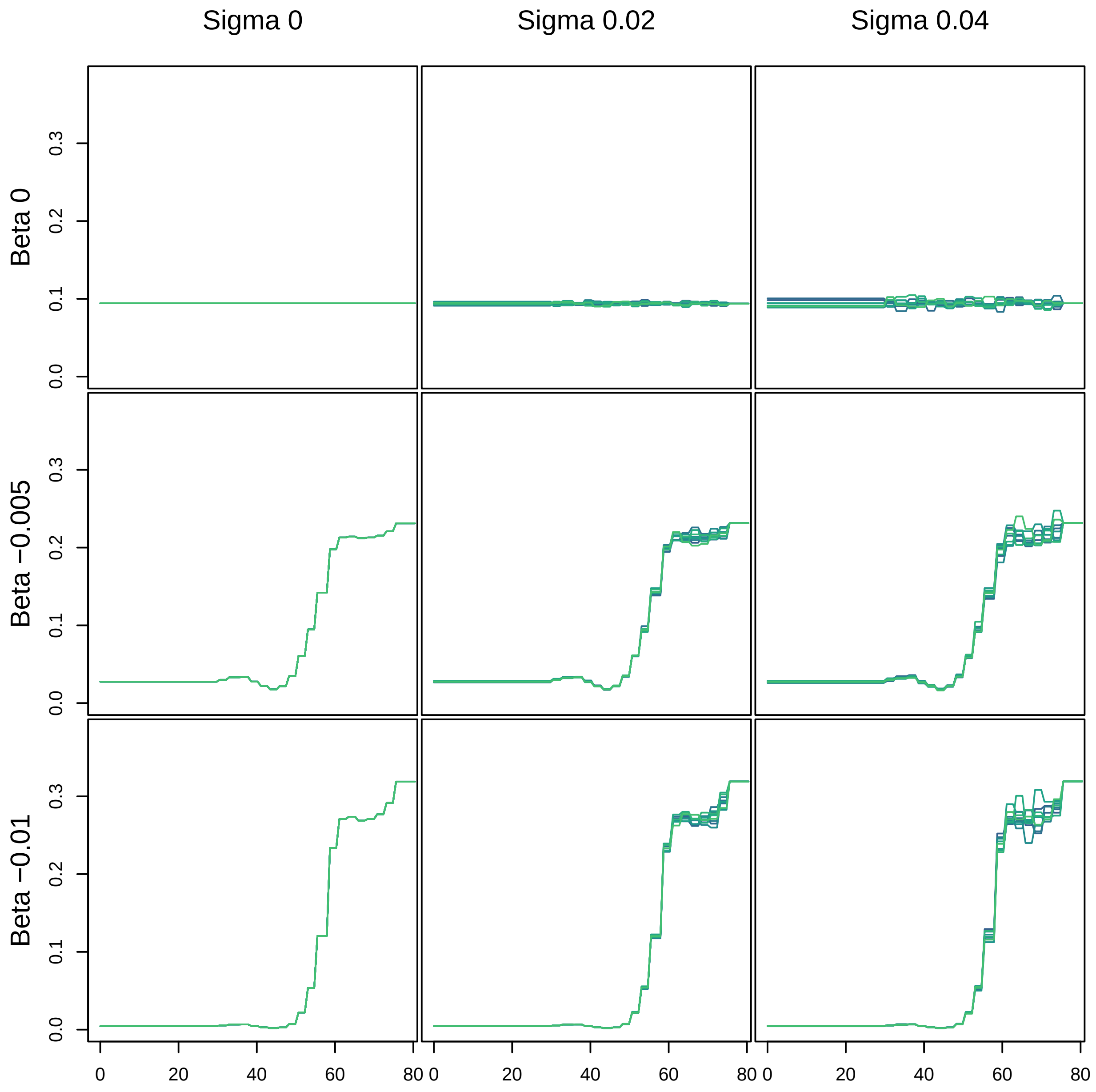

**Figure S16: Speciation rated simulated under an environmentally-dependent UC diversification model.** We simulated speciation rates with different correlation coefficients $\beta = \{0, -0.005, -.01\}$ and standard deviations $\sigma = \{0, 0.02, 0.04\}$. We used the atmospheric $CO_2$ as the environmental variable. When $\beta = 0$ in the simulations, then this model was equivalent to an environmentally-independent diversification model. When $\sigma = 0$ in the simulations, then this model was equal to the *fixed* environmentally-diversification model. When both $\beta = 0$ and $\sigma = 0$, then we simulated under a constant rate process (top left). This establishes the false positive rate in our simulations. Note that the autocorrelated model (GMRF, Figure S15) has overall more variation in speciation rates (e.g., top right) compared to the uncorrelated model because the variation is additive over epochs in the autocorrelated model.

#### S18.2 Simulated phylogenies under environmentally-dependent diversification rates

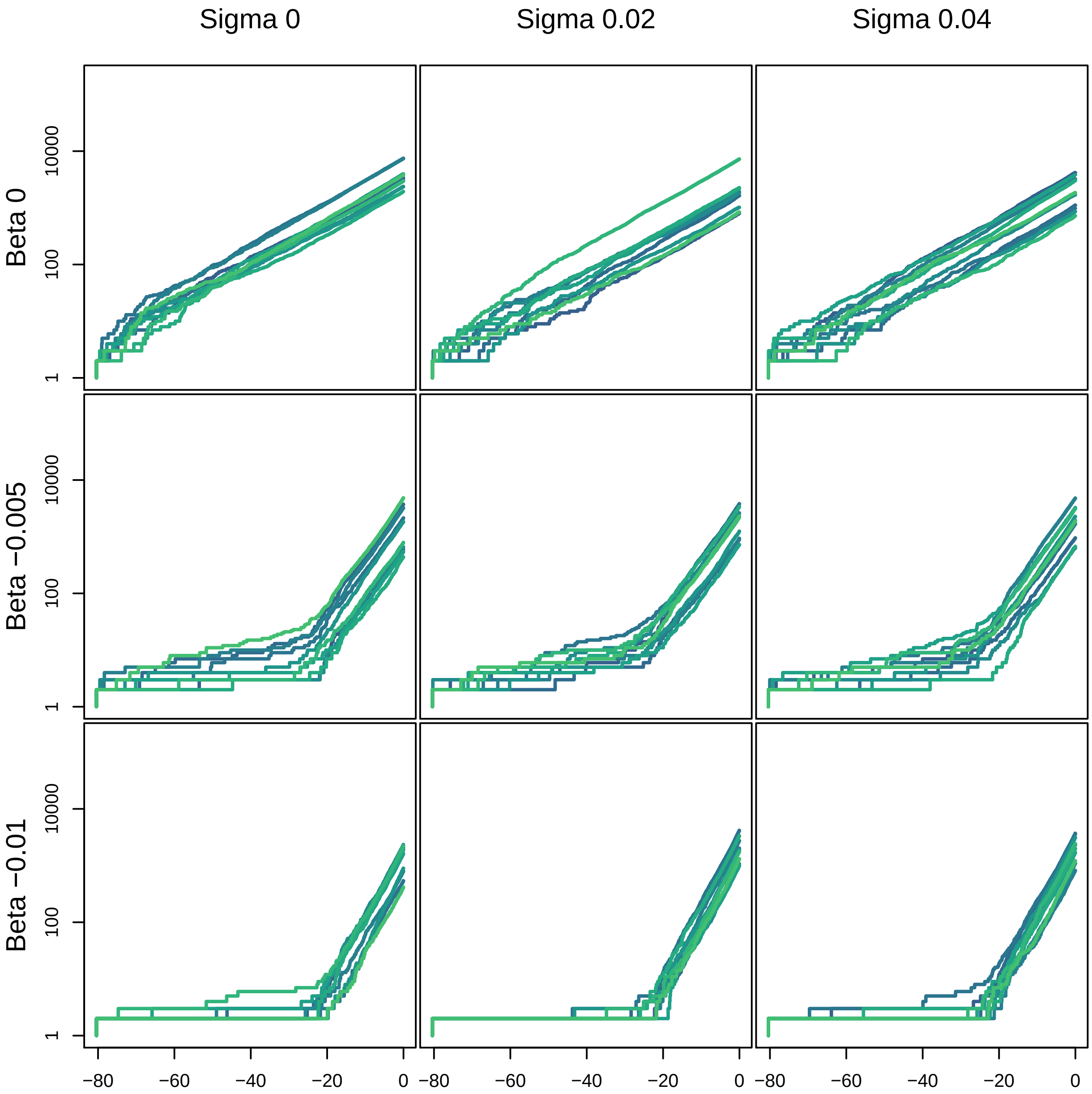

**Figure S17: Lineage-through-time (LTT) curves of 10 phylogenies simulated under an environmentally-dependent GRMF diversification model.** We used the simulated diversification rate trajectories shown and described in Figure S15. We nicely observe the log-linear LTT curve for the constant-rate birth-death simulations (top left) and increasing deviation from the log-linear curve with higher correlation coefficient $\beta$. Higher standard deviation $\sigma$ in the speciation rates led to slightly more variation in LTT curves but no large effect.

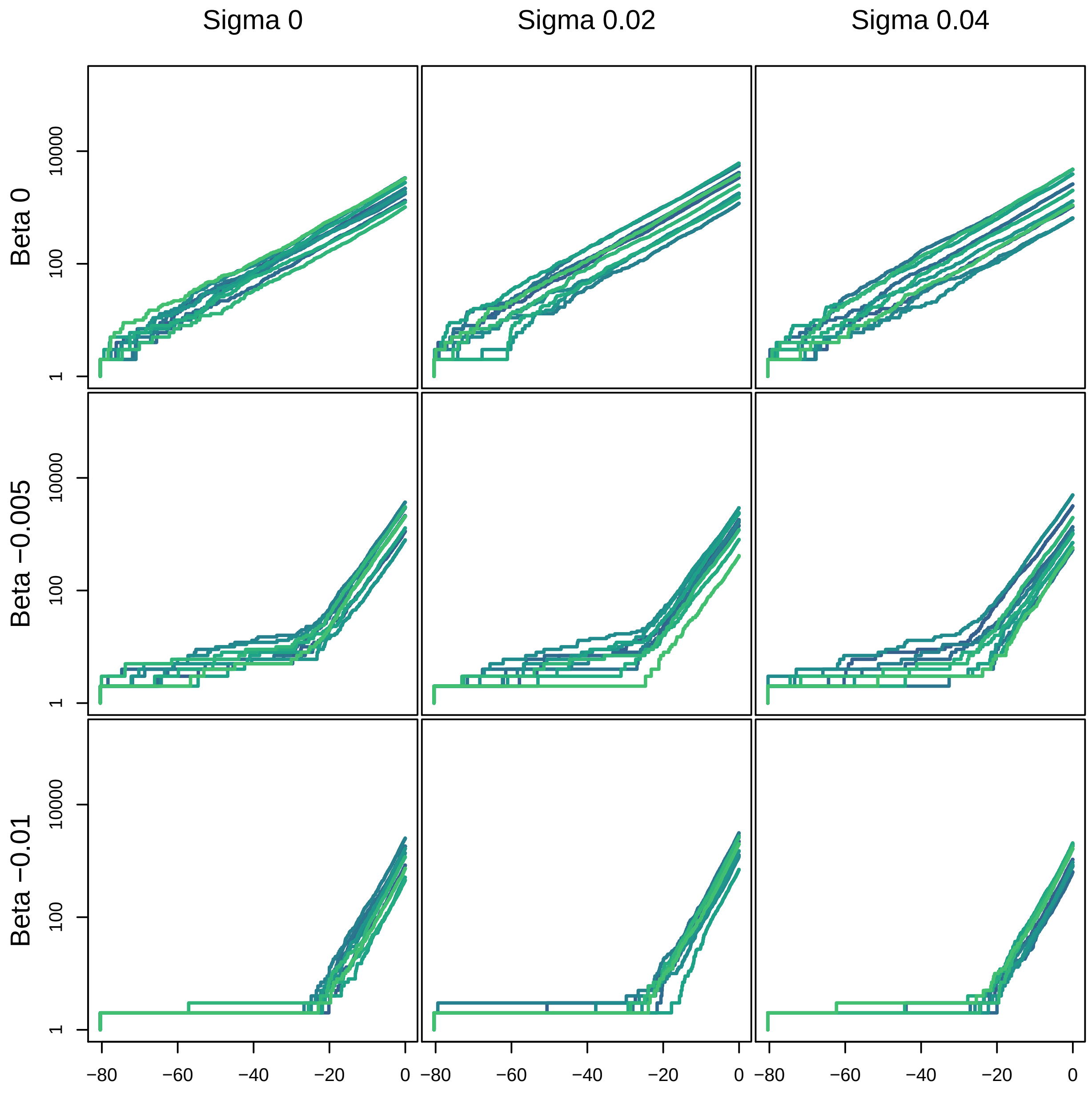

**Figure S18: Lineage-through-time (LTT) curves of 10 phylogenies simulated under an environmentally-dependent UC diversification model.** We used the simulated diversification rate trajectories shown and described in Figure S16. We nicely observe the log-linear LTT curve for the constant-rate birth-death simulations (top left) and increasing deviation from the log-linear curve with higher correlation coefficient $\beta$. Higher standard deviation $\sigma$ in the speciation rates led to slightly more variation in LTT curves but no large effect.

##### S18.3 Estimated correlation coefficients from the phylogenies simulated under environmentally-dependent diversification rates

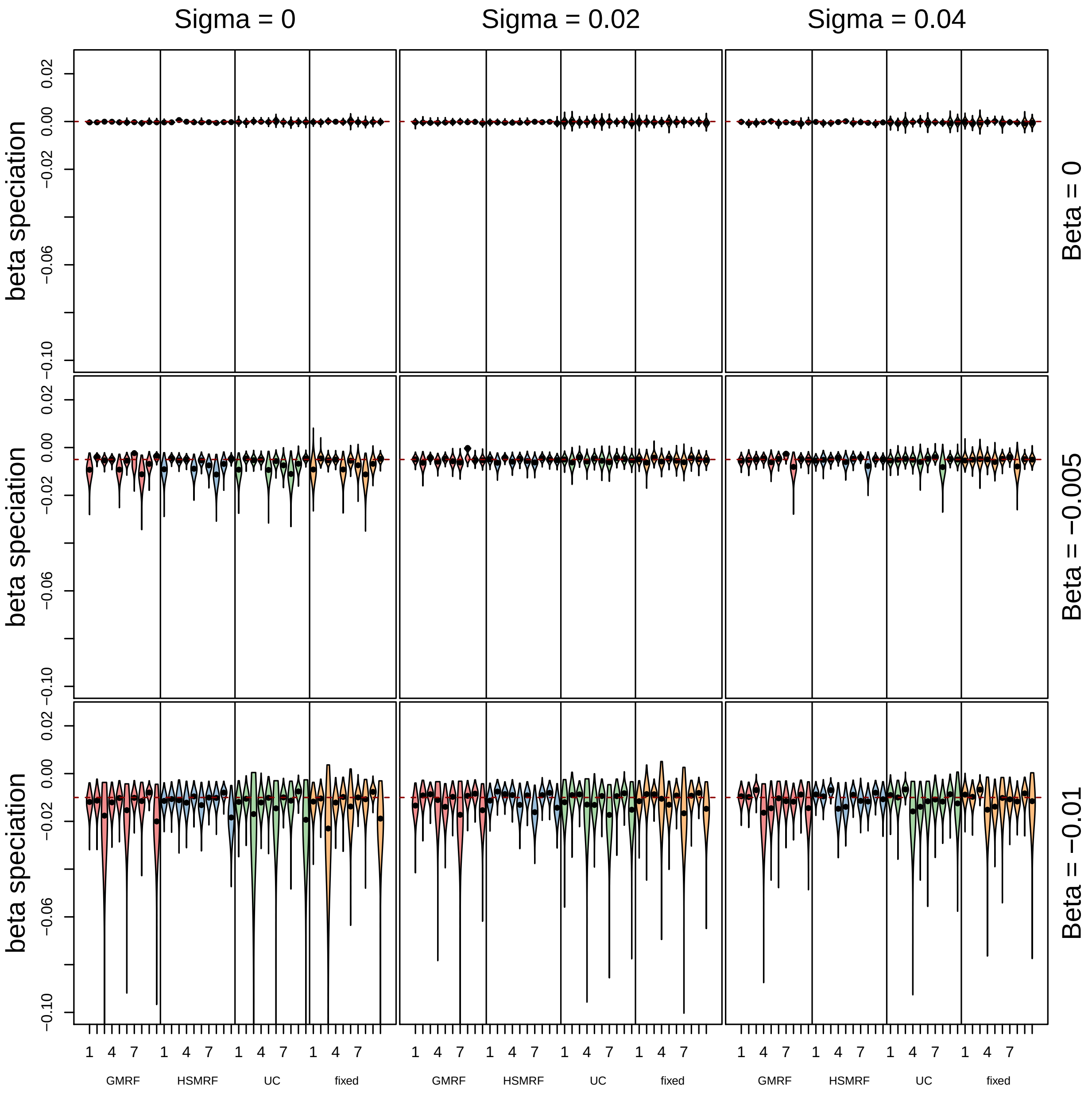

**Figure S19: Estimated correlation coefficients for the simulated phylogenies under the GMRF environmentally-dependent diversification rates model.** We estimated correlation coefficient $\beta$ using the simulated phylogenies shown and described in Figure S17. We used all four environmentally-dependent diversification rates model for inference, running in total 450 MCMC analyses. Overall, all four models perform well to estimate the environmental correlation. As we noticed in our simulations, the diversification rate variation is primarily driven by the environmental variable and less by additional factors (comparably low $\sigma$). In these situations it does not matter much which environmentally-dependent diversification rates model is used. Nevertheless, the HSRMF model always produced the smallest credible intervals and was most precise. This simulation study shows that there is high power to detect environmental correlation to diversification rates (middle and bottom row) while retaining a low false-positive rate (top row). The results are congruent with our inferences using the simulated trees under the UC environmentally-dependent diversification rates model (Figure **??**).

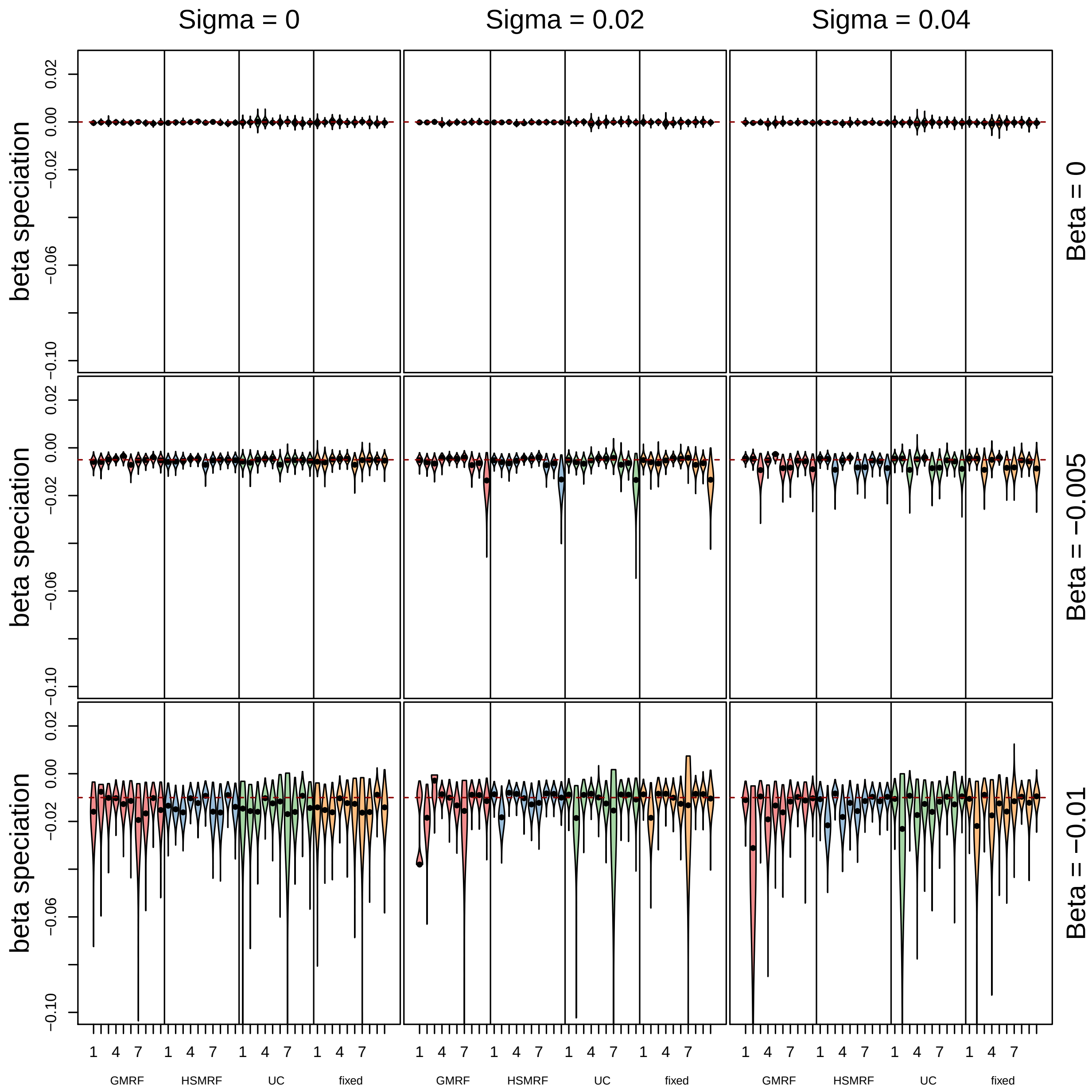

**Figure S20: Estimated correlation coefficients for the simulated phylogenies under the UC environmentally-dependent diversification rates model.** We estimated correlation coefficient $\beta$ using the simulated phylogenies shown and described in Figure S18. We used all four environmentally-dependent diversification rates model for inference, running in total 450 MCMC analyses. Overall, all four models perform well to estimate the environmental correlation. As we noticed in our simulations, the diversification rate variation is primarily driven by the environmental variable and less by additional factors (comparably low $\sigma$). In these situations it does not matter much which environmentally-dependent diversification rates model is used. Nevertheless, the HSRMF model always produced the smallest credible intervals and was most precise. This simulation study shows that there is high power to detect environmental correlation to diversification rates (middle and bottom row) while retaining a low false-positive rate (top row). The results are congruent with our inferences using the simulated trees under the GMRF environmentally-dependent diversification rates model (Figure S20).

#### S19 Simulation study using empirical taxon sampling

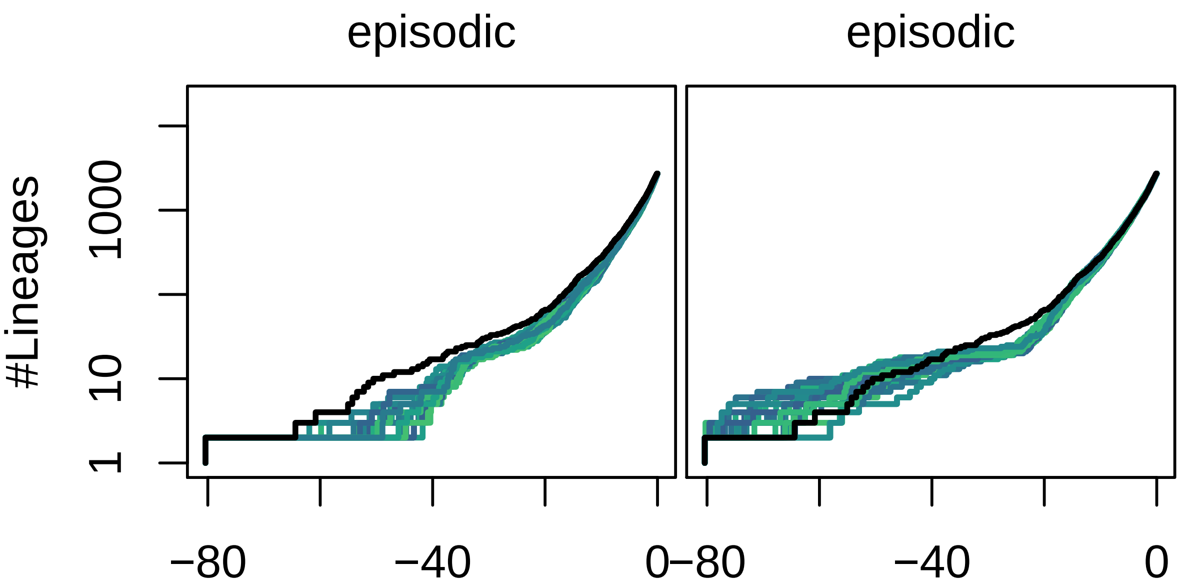

**Figure S21: Lineage-through-time curves of simulated phylogenies under empirical sampling.** We simulated 100 phylogenies (only first 10 shown here) under a constant-rate birth-death process (false-positive) and an episodic-birth-death process (power) using empirical taxon sampling. These trees were simulated by adding the missing species randomly in the pre-assigned clades of the daisy phylogeny. Then, we simulated divergence times under either the constant or episodic birth-death process. After simulation, the additional taxa were pruned again to mimic empirical sampling. The solid black line shows the empirical LTT curve of the daisy phylogeny.

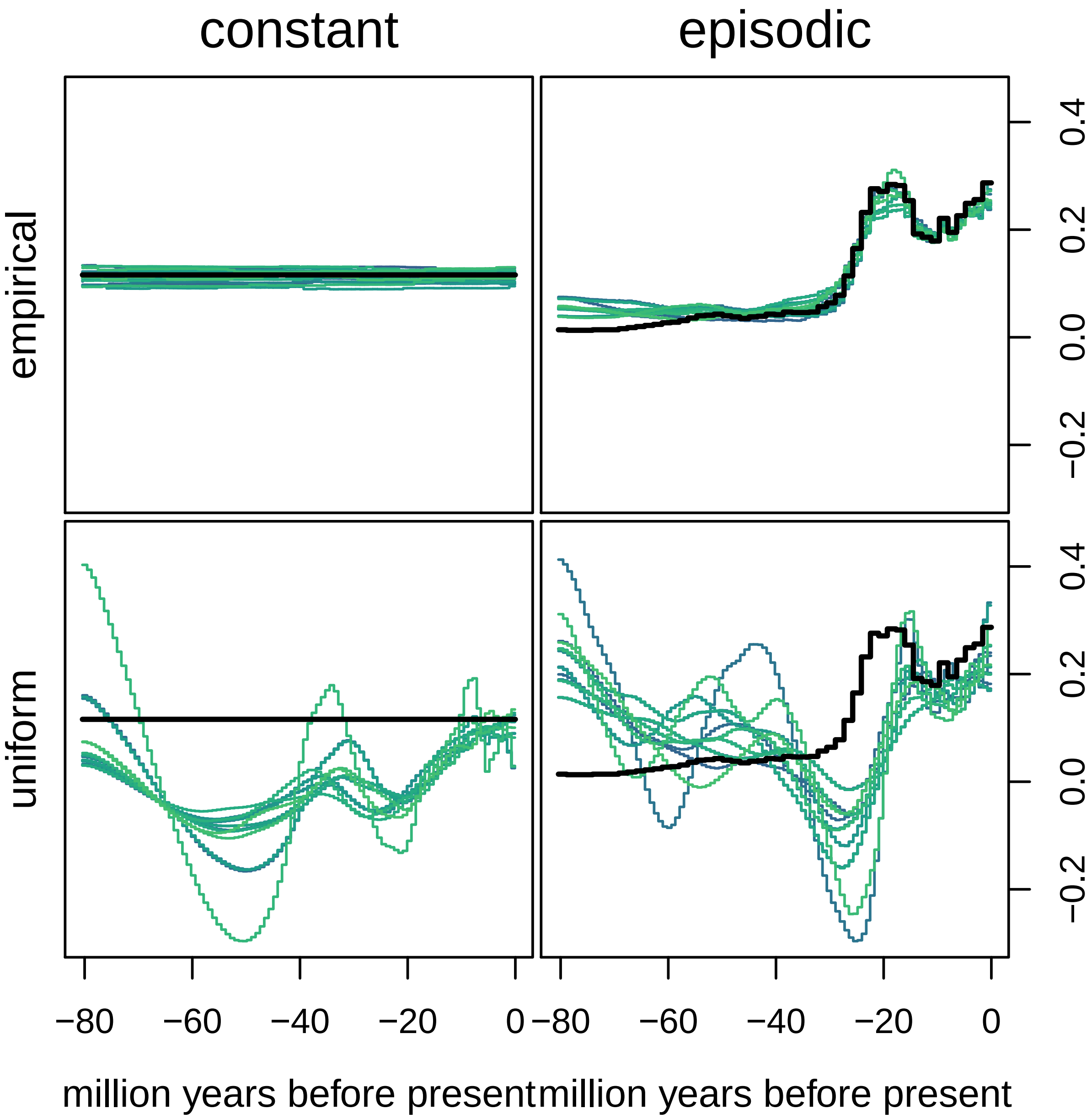

**Figure S22: Estimated net-diversification rates using the simulated phylogenies from Figure S21**. The solid black line shows our true net-diversification rates. If our assumption of incomplete taxon sampling matches the simulation conditions, then we infer unbiased diversification rates, as previously shown under similar incomplete taxon sampling schemes [11, 10, 17]. Conversely, if the incomplete taxon sampling assumption is violated, e.g., assuming all missing species are *uniformly* distributed within the phylogeny, then we obtain strongly biased diversification rates. We also note that our GMRF model shows a good performance even when the true diversification rates are constant (top left, low false-positive rate). Our good precision in not surprising because the simulated phylogenies had 2723 tips, as our daisy phylogeny.
